# Neuromodulation of *Foxp2*^+^ hypothalamic neurons induces therapeutic hypothermia

**DOI:** 10.64898/2026.05.04.722579

**Authors:** Guangsen Bao, Ling Zhang, Mei Li, Lu Wang, Ting Yu, Zewen Zhu, Jinghan Cui, Yunmei Zhu, Xiaofeng Lin, Jiayi Cong, Guoyuan Liu, Dengke K. Ma, Zhe Zhang, Mingliang Ye, Biao Yu, Xiaheng Zhang, Lei Xiao, Wei Jiang, Yongjun Dang

**Author notes:** These authors contributed equally: Guangsen Bao and Ling Zhang.

## Abstract

The inability to selectively trigger therapeutic hypothermia independent of environmental cooling has hindered causal analysis of its broader physiological benefits. Here, we employed P57, a natural product that pharmacologically induces therapeutic hypothermia circumventing external cold stress, to identify the neuronal substrates underlying hypothermia-induced antitumor effects. Integrating functional ultrasound imaging, activity-dependent neuronal labeling, and single-nucleus RNA sequencing, we identify a previously unrecognized population of *Foxp2*⁺ neurons in the medial preoptic area (MPA) that are selectively activated by P57. Inhibition of MPA *Foxp2^+^* neurons abolishes P57-induced hypothermia, whereas the paraventricular hypothalamus (PVH) serves as a critical downstream node. Importantly, targeted activation of MPA *Foxp2*⁺ neurons is sufficient to induce sustained hypothermia, suppress systemic metabolism, and inhibit tumor growth, providing causal evidence that neuronally driven reductions in core body temperature can exert antitumor effects. Together, these findings establish MPA *Foxp2*⁺ neurons as a controllable node for therapeutic hypothermia induction and demonstrate that neuromodulation of defined neuronal populations can achieve therapeutic benefit by directly controlling physiological states.

---

Temperature is a fundamental biophysical variable that shapes biological processes across molecular, cellular, and systemic scales, ultimately influencing physiology, behavior, survival, and disease susceptibility, whether through environmental exposure or changes in core body temperature^1^. For instance, long-lived bowhead whales inhabiting Arctic waters exhibit enhanced DNA repair^2^, hibernating brown bears maintain low body temperature and are protected from venous thromboembolism^3^, and hypothermic torpor in laboratory mice markedly slows epigenetic aging^4^. Reduced ambient or core temperature has been associated with beneficial effects in cancer, obesity, chronic inflammatory disorders, and age-related physiological decline^5–9^. Clinically, hypothermia has also long been applied from ancient cold compresses to modern cardiac surgery^10,11^. However, adaptive thermogenic and metabolic responses triggered by environmental cold exposure are fundamentally distinct from the systemic metabolic suppression associated with true hypothermia. Environmental temperature changes evoke strong compensatory thermoregulation, complicating attribution of physiological effects specifically to hypothermia. Besides, conventional physical cooling approaches suffer from uneven temperature control, local cold injury, and limited precision, restricting their broader clinical applicability. These observations raise two central questions motivating our work: does hypothermia itself directly contribute to health promotion and disease intervention, and whether hypothermia can be induced in a safer and more precisely controlled manner than conventional physical cooling thereby unlocking its substantial therapeutic potential.

Understanding how the brain controls body temperature offers a path to overcome the limitations of current cooling methods. Recent studies have identified distinct populations of thermoregulatory neurons, providing an unprecedented view into central control of temperature. Different stimuli activate different subsets of these neurons, such as those regulating torpor^12^, sensing light^13,14^, expressing neuropeptides^15^, responding to hormonal signals^16^, or reacting to cold and heat^17–19^, resulting in stimulus-specific patterns of temperature change^20^. These findings highlight the heterogeneity of thermoregulatory neurons and raise the possibility that certain neuronal populations may be optimally suited for inducing a therapeutically beneficial hypothermic state due to their unique downstream effectors. Following this rationale, we previously found that the natural product P57 induces hypothermia by inhibiting Pdxk, a key kinase in vitamin B6 metabolism, thereby increasing the accumulation of its substrate, pyridoxal (PL), in the hypothalamus. P57-induced hypothermia protects against nerve injury in rodent middle cerebral artery occlusion models without detectable systemic toxicity^21^, suggesting its potential as a tool to explore neuron-driven therapeutic hypothermia. Here, we identify a population of *Foxp2*^+^ neurons in the medial preoptic area (MPA) as key regulators of P57-induced hypothermia, and show that targeted neuromodulation of these neurons provides a safe and controllable strategy for tumor suppression, directly demonstrating the therapeutic benefit of hypothermia. This evidence further illustrates how neuromodulation of physiological states can achieve therapeutic benefit, extending its potential beyond traditional neurological and psychiatric disorders.

## Results

### P57 activates neurons in MPA and PVH

We previously showed that P57 induces hypothermia via a pharmacological mechanism involving altered vitamin B6 metabolism^21^. However, its effect on the central nervous system, especially on neuronal activity in the thermoregulatory center of the hypothalamus, remains unclear. To address this, we used functional ultrasound (FUS) imaging, which captures brain hemodynamic activity based on neurovascular coupling, to reflect real-time activity changes across brain regions during P57-induced hypothermia by calculating the change of average relative cerebral blood volume compared to baseline and control (ΔrCBV)^22^. Given that anesthetics can substantially alter cerebral blood volume (CBV), we performed FUS imaging in awake, head-fixed mice after habituation to minimize stress-related effects (Fig. S1a). Following intraperitoneal injection (i.p.) of P57, rapid brain-wide changes of CBV were observed. Within 5 minutes, 44.5% of the imaged brain regions exhibited a robust rise in ΔrCBV, indicating elevated neuronal activity (Fig. S1b and c). Projection of these activity maps onto the Allen Mouse Brain Atlas revealed sustained activation predominantly in the isocortex, thalamus, and hypothalamus (Fig. 1a), which are associated with somatosensory^23^, sensory transmission^24^, and body temperature regulation^25^, respectively. Subsequently, ΔrCBV declined progressively, and by 40 min post-injection, activity in all imaged regions was suppressed (Fig. S1b). This global suppression paralleled the progressive drop in body temperature, and the rectal temperature (T_R_) of mice fell below 34 ℃ at 40 min after P57 administration^21^, indicating a moderate hypothermic state that leads to hypovolemia^26^. Specifically, within the hypothalamus, the median eminence (ME) exhibited robust and sustained activation, lasting nearly 30 minutes. The hypothalamic medial zone (MEZ), which includes the core thermoregulatory region of the preoptic area (POA)^27^ and the paraventricular hypothalamic nucleus (PVH), a region implicated in thermoregulatory circuits^28–30^, was also activated following P57 treatment (Fig. 1b and c, Fig. S1d-f). These results reveal real-time, region-specific neural dynamics during P57-induced hypothermia.

**Fig. 1.**
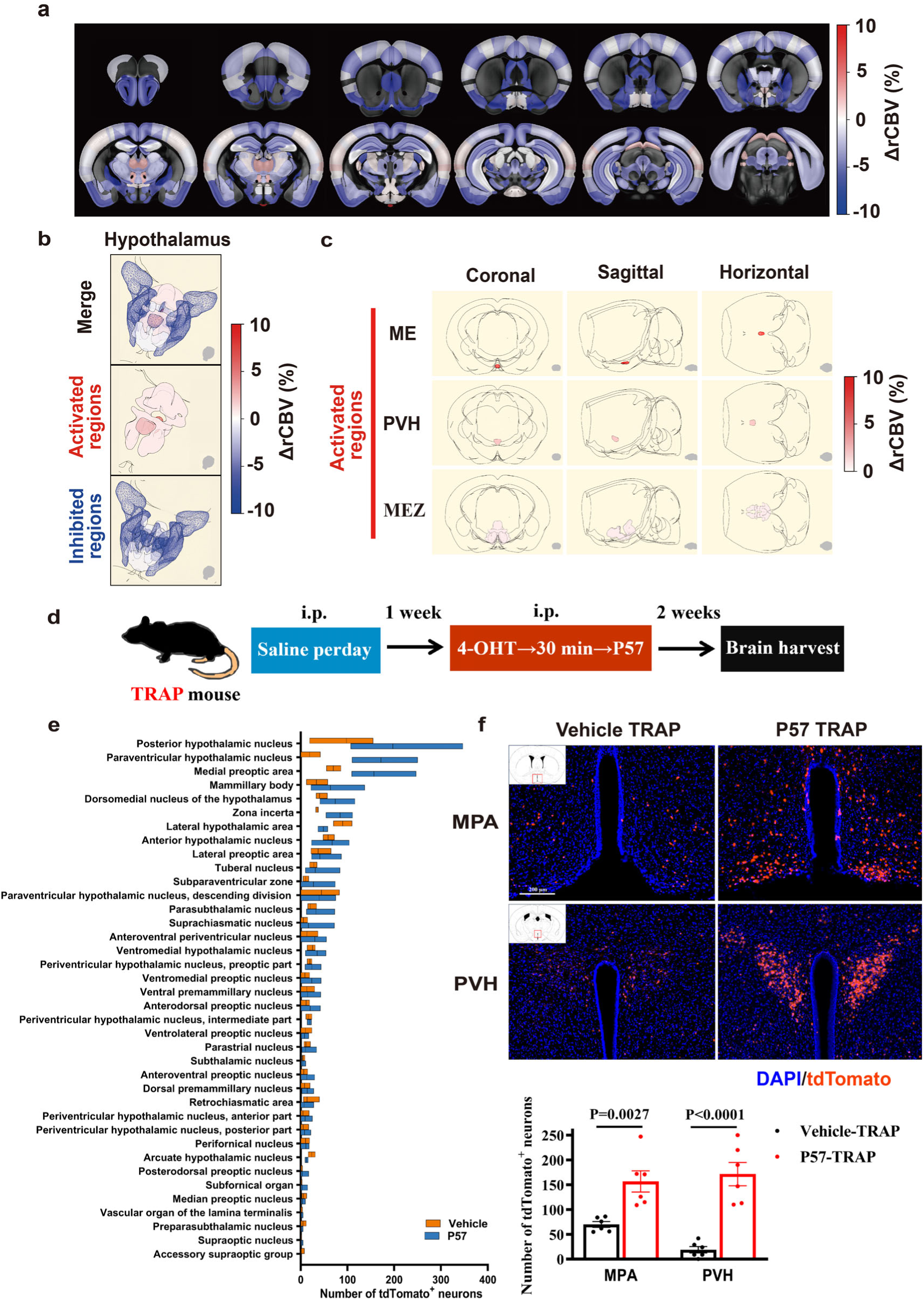
Real-time functional mapping and neuronal activity tracking identify MPA and PVH as a key area for P57-induced hypothermia. **a**, Average relative cerebral blood volume changes (ΔrCBV) of different region across the brain within 5 to 10 minutes after administration P57 compared to vehicle. **b,** 3D rendering of ΔrCBV of different hypothalamic nuclei within 5 to 10 minutes after administration P57 compared to vehicle. **c,** Different views of the 3D rendering results of various P57-activated brain regions in the hypothalamus within 5 to 10 minutes after administration. **d,** Schematic of labeling P57 activated neurons using *Fos-CreERT2;tdTomato* mice (n=6 mice for each group). **e, f,** Number of tdTomato^+^ neurons across different hypothalamic nuclei **(e)** and representative slice (top) and quantification (bottom) of MPA and PVH **(f)** in P57 group and Vehicle group in *Fos-CreERT2;tdTomato* mice. Student’s t test used was two-sided. Heat-map values are means; floating bars show min to max and mean (line). Bar chart and error bars are presented as mean values ± s.e.m. Median eminence (ME), Paraventricular hypothalamic nucleus (PVH), Hypothalamic medial zone (MEZ),

To translate this system-level insight into a neuron-type-targeted strategy for therapeutic hypothermia, we next refined our analysis to single-neuron resolution. *Fos-CreERT2;tdTomato* mice (hereafter referred to as TRAP mice), specifically developed for activity-dependent neuronal labeling^31^, were used to mark neurons activated by P57 that may possess hypothermic induction potential (Fig. 1d). TRAP mice express a 4-hydroxytamoxifen (4-OHT)-dependent CreERT2 recombinase under the *Fos* promoter. Consequently, CreERT2 is transiently synthesized in activated neurons and is functional only in the presence of 4-OHT. The recombinase excises a transcriptional stop cassette upstream of tdTomato, driving permanent red fluorescence to precisely mark neurons activated by a specific stimulus. When TRAP mice received both 4-OHT and P57, only neurons activated by P57 were labelled with tdTomato. By quantification of tdTomato-positive cells across hypothalamic subregions, we observed results that were relatively consistent with the FUS mapping data, with specific hypothalamic regions being activated (Fig. 1e, Fig. S2a). Among these, tdTomato-positive cells were significantly increased in the MPA and PVH, the core regions of thermoregulation (Fig. 1f). These data demonstrate that neurons in MPA and PVH are specifically activated in response to P57 treatment.

### Molecular characteristics of P57-activated MPA neurons

While TRAP labeling shows that P57 selectively activates MPA and PVH neurons, direct evidence that these labeled neurons can induce hypothermia is lacking. Moreover, previous work found that direct injection of P57 into the PVH fails to lower body temperature^21^, prompting us to focus on the P57-activated population within the MPA. To directly test whether these neurons are sufficient to induce hypothermia, we utilized Gq-DREADD (Gq-coupled designer receptors exclusively activated by designer drug) in *Fos-CreERT2* mice. This strategy enabled CreERT2-dependent expression of DIO-Gq-mCherry specifically in P57-activated neurons (Fig. 2a and b). Administration of clozapine N-oxide (CNO, the DREADD agonist) to activate these MPA neurons resulted in a significant reduction in body temperature (Fig. 2c). These findings demonstrate that P57-activated MPA neurons are the key neurons for P57-induced hypothermia. Thus, these neurons represent a promising neuronal target for therapeutic hypothermia.

**Fig. 2.**
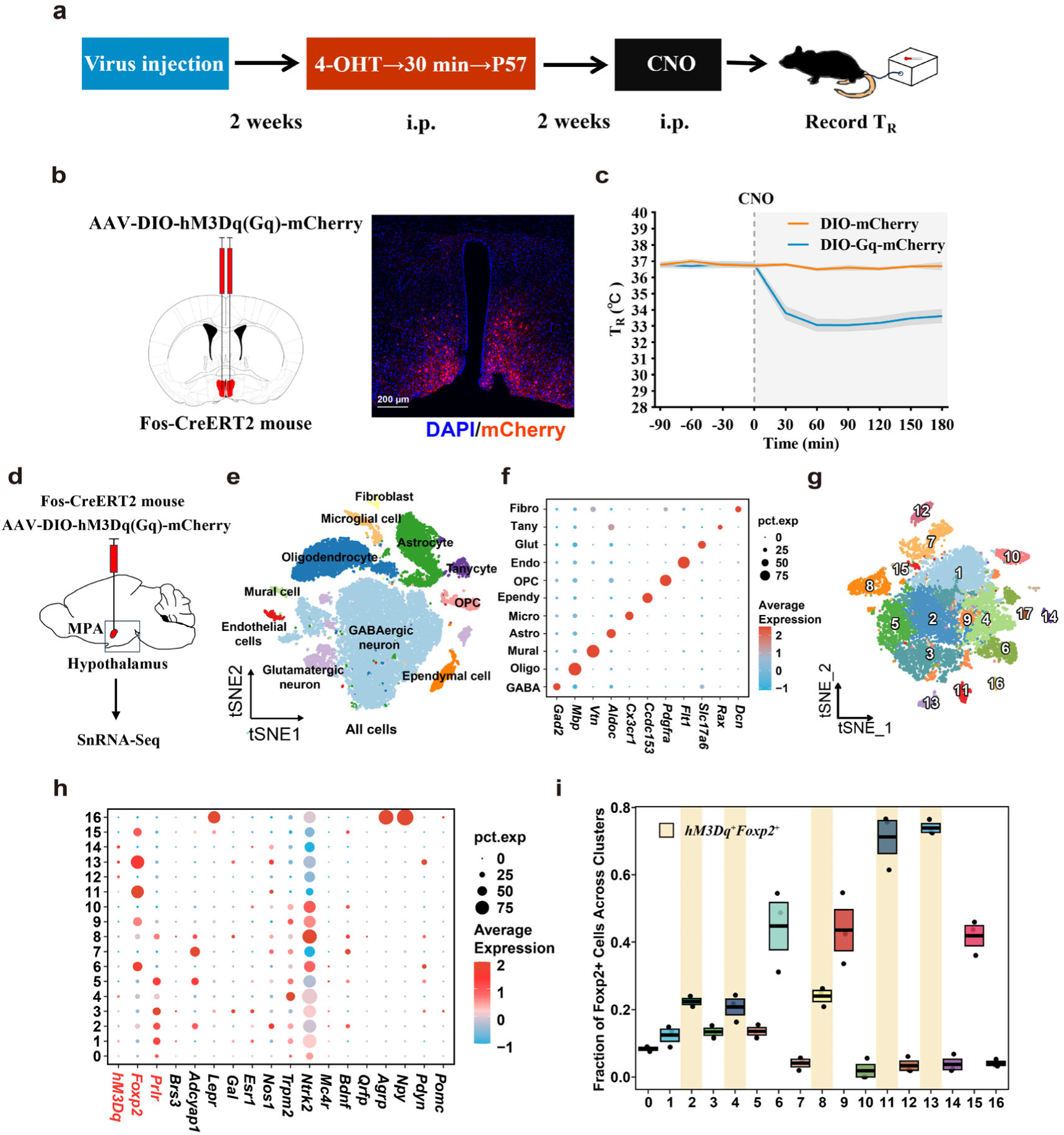
Molecular characterization of P57-activated neurons that induced hypothermia. **a,** Schematic shows the process of using chemogenetics to manipulate neurons specifically activated by P57 and detect body temperature in *Fos-CreERT2* mice. **b,** Schematic diagram of the AAV-DIO-hM3Dq(Gq)-mCherry (DIO-Gq-mCherry) injection site (left) and the expression of the virus in MPA in *Fos-CreERT2* mice (right). **c,** Rectal temperature (T_R_) of *Fos-CreERT2* mice, that received virus injection (n=8 mice in DIO-Gq-mCherry group and 6 mice in DIO-mCherry group) and were induced to express by 4-OHT and P57, before and after intraperitoneal injection of CNO. The dashed line indicates the onset of CNO administration, shading indicates error bars. **d,** Schematic showing the procedure for the molecular characterization of P57-activated neurons that induced hypothermia. AAV-DIO-hM3Dq(Gq)-mCherry is injected into the MPA (n = 3 mice). After induced expression, the hypothalamus is dissected and analyzed by snRNA-seq. **e,** Uniform manifold approximation and projection (UMAP) plot of 26,087 nuclei from the MPA of 3 mice. Colors distinguish the main cell types. OPCs, oligodendrocyte precursor cells. **f,** Expression of the indicated marker genes across different cell types (named as abbreviations of the cell types in **(e)**). **g,** UMAP plot of 15,328 neuronal nuclei. Colors distinguish the 17 neuronal subtypes. **h,** Expression of *hM3Dq* and the marker genes of temperature-sensing or regulating neurons reported previously in various clusters. **i,** Distribution of *Foxp2*^+^ neurons across all neuronal cell types. For the box plots, the centre line and box boundaries indicate mean ± s.e.m. (n = 3 mice). Yellow shading indicates cell types expressing *Foxp2* and *hM3Dq*.

Given the extensive heterogeneity of hypothalamic neuron types in both mice and humans, even within the MPA^32,33^, we next sought to define the molecular identity of this functionally defined neuronal subtype. We performed single-nucleus RNA-sequencing (snRNA-seq) on hypothalamic cells, including P57-activated, hM3Dq-positive neurons with hypothermia-inducing function. Hypothalamus from three mice that had been both TRAP-labelled and functionally verified to drive hypothermia were dissociated for snRNA-seq (Fig. 2d). A total of 26,087 high-quality single nuclei were profiled. Following dimensionality reduction, clustering, and annotation, we resolved the major hypothalamic cell classes (Fig. 2e and f, Fig. S3a), comprising 15,328 neurons. Among these, 646 neurons expressed *hM3Dq*, identifying them as the P57-activated, hypothermia-inducible population (TRAPed neurons) (Fig. S3b). To further characterize this population, we subdivided the 15,328 neurons into 17 subclusters (Fig. 2g and Fig. S3c-e) and examined each for previously reported thermoregulatory marker genes (both hypothalamic and extra-hypothalamic). Subclusters with high proportion and expression level of *hM3Dq* also exhibited relatively high frequency and expression of *Foxp2* (Fig. 2h, i, and Fig. S4a-c), implicating *Foxp2*-rich subclusters as components of the P57-responsive ensemble. In parallel, we used *hM3Dq* expression itself as the grouping criterion, separating all neurons into *hM3Dq-positive* (*hM3Dq*⁺) and *hM3Dq*-negative (*hM3Dq*⁻) populations (Fig. S4d-f). *Prlr* was selectively enriched in *hM3Dq*⁺ neurons (Fig. S4g–i), and inspection of the 17 subclusters revealed that *Prlr* and *Foxp2* exhibit distinct, non-overlapping distribution preferences. These results indicate that *Foxp2* and *Prlr* may serve as molecular markers for P57-activated MPA neurons.

### *Foxp2*^MPA^ neurons are necessary for P57-induced hypothermia

Transcriptomic analysis identified neuronal subsets in the MPA expressing either *Foxp2* or *Prlr* (*Foxp2*^MPA^ and *Prlr*^MPA^ neurons) as candidate neurons of P57-induced hypothermia. To validate this finding, we performed immunofluorescence on brain slices from labeled *Fos-CreERT2* mice. Co-staining revealed that among P57-activated neurons in the MPA, 37.93 ± 1.93% (mean ± s.e.m) expressed Foxp2, 82.97 ± 1.25% expressed Prlr, and 30.77 ± 1.69% expressed both markers (Fig. 3a and b). Consistent with these results, co-labeling of Foxp2 (32.35 ± 2.61%), Prlr (60.66 ± 3.10%), or Foxp2/Prlr (23.78 ± 1.65%) with c-Fos protein was also found in MPA in C57BL/6 mice treated with P57 (Fig. 3c and d). Moreover, we also observed the expression specificity of *Foxp2* and *Prlr* in MPA in the Allen mouse brain atlas (Fig. S5a). Notably, one recent study reported that selective activation of *Prlr-*positive neurons in POA is sufficient to drive deep hypothermia (core body temperature <30℃)^34^. This convergence independently validates both our labeling strategy and our transcriptomic identification.

**Fig. 3.**
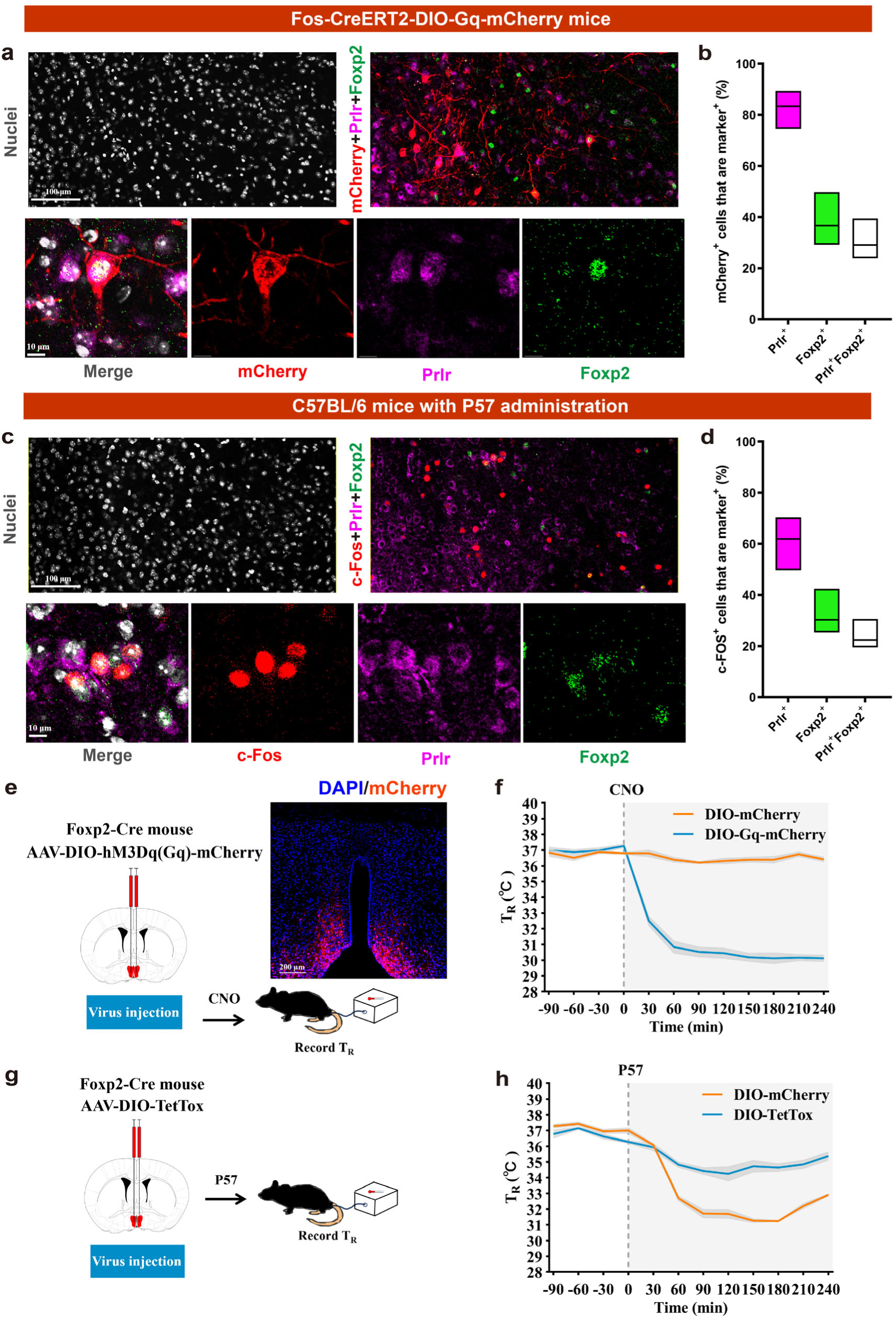
Activating *Foxp2*^MPA^ neurons lowers body temperature and is necessary for P57-induced hypothermia. **a, b,** Low-magnification and magnification images of nuclei (DAPI) and mCherry, Foxp2, PRLR expression in MAP in *Fos-CreERT2* mice injected AAV-DIO-hM3Dq(Gq)-mCherry virus **(a)** with quantification (n=11 mice), floating bars show min to max and mean (line) **(b)**. **c, d,** Low-magnification and magnification images of nuclei (DAPI) and c-Fos (red, pseudo color), Foxp2 (green, pseudo color), PRLR (magenta) expression in MAP in C57BL/6 mice intraperitoneal injection of P57 after 2 h **(c)** with quantification (n=6 mice) **(d)**. **e,** Schematic diagram of the AAV-DIO-hM3Dq(Gq)-mCherry injection site (left) and the expression of the virus in MPA in *Foxp2-Cre* mice (right). **f,** The T_R_ of *Foxp2-Cre* mice that received virus injection (n=9 mice in DIO-Gq-mCherry group and 5 mice in DIO-mCherry group) before and after intraperitoneal injection of CNO. The dashed line indicates the onset of CNO administration, shading indicates error bars. **g,** Schematic diagram of the AAV-DIO-TetTox injection in MPA in *Foxp2-Cre* mice. **h,** The T_R_ of Foxp2-Cre mice that received virus injection (n=4 mice in DIO-TetTox group and 4 mice in DIO-mCherry group) before and after intraperitoneal injection of P57. The dashed line indicates the onset of CNO administration, shading indicates error bars.

On the basis that the hypothermia-inducing capacity of *Prlr*-positive neurons has now been established, we next focused on determining the functional contribution of the *Foxp2*-positive subpopulation. We injected the DIO-hM3Dq-mCherry virus into the MPA of *Foxp2-Cre* mice. Selective chemogenetic activation of *Foxp2*^MPA^ neurons was sufficient to markedly lower core body temperature (T_R_<30℃) (Fig. 3e and f). Furthermore, expressing Cre-dependent Tetanus toxin light chain (DIO-TetTox) in *Foxp2*^MPA^ neurons to abolish synaptic transmission, the P57-induced hypothermia was blocked (Fig. 3g and h). Taken together, these results demonstrate that *Foxp2*^MPA^ neurons are both necessary and sufficient for P57-induced hypothermia.

### PL activates the ERK-FOS pathway

Our study characterized the molecular identity of P57-activated neurons in MPA and demonstrated both their sufficiency to induce hypothermia and their necessity for P57-induced hypothermia. We next sought to identify the factor that mediates the selective activation of these cells by P57. Considering that P57 induced hypothermia by inhibiting Pdxk kinase activity and leading to PL accumulation in the hypothalamus^21^, we hypothesized that the regional expression pattern of Pdxk might determine neuronal responsiveness. Single-cell transcriptome analysis revealed that *Pdxk* was significantly enriched in P57-activated neurons (*hM3Dq*^+^ neurons) relative to P57-unresponsive neurons (*hM3Dq*^-^ neurons) (Fig. S6a). However, co-immunostaining showed that only a very small number of c-Fos-positive cells express Pdxk (4.22 ± 0.38%) in MPA after P57 administration (Fig. S6b). Given the relatively low proportion of Pdxk-expressing neurons within this population, Pdxk expression alone is unlikely to account for their selective activation by P57. This led us to postulate that P57-activated neurons may possess an unidentified PL target protein, which triggers neuronal excitation.

To test this hypothesis, we employed Peptide-centric Local Stability Assay (PELSA)^35^ to identify PL-binding proteins in the hypothalamus, and integrated these data with differentially expressed genes (DEGs) analysis comparing *hM3Dq*^+^ and *hM3Dq*^-^neurons. Intersection of the PL-binding proteins with the top 100 DEGs yielded a single candidate: Acadsb (Fig. S6c). PELSA assays confirmed that PL binding increases Acadsb stability (Fig. S6d), and snRNA-seq data revealed Acadsb enrichment in *hM3Dq*⁺ neurons (Figure. S6e). We verified the PL–Acadsb interaction by thermal shift assay, and confirmed the co-labeling of Acadsb and c-Fos (39.29 ± 1.71%) (Fig. S6f and g). These findings suggest that TRAPed neurons may be selectively excited by PL, potentially due to their enriched expression of Acadsb, a previously unrecognized PL target protein.

We next investigated the mechanism by which PL activates these neurons via Acadsb. Affinity-capture mass spectrometry data from the Biological General Repository for Interaction Datasets (BioGRID) identified RAF1 as an Acadsb-interacting protein^36^. Given that the RAF1-MEK-ERK pathway is the classic pathway for initiating *FOS* gene transcription^37–39^, we examined phosphorylation of its terminal effector ERK in the MPA 30 min after intraperitoneal injection of PL. Compared with the vehicle-treated controls (29.00 ± 1.63%), PL elevated the proportion of pERK1/2 (T202/Y204)-positive cell number among Acadsb-positive cells to 44.00 ± 1.80% (Fig. S6h and i). These findings support a model in which PL engages the ERK-FOS pathway to activate MPA neurons, potentially acting through Acadsb.

### P57-activated MPA neurons project to PVH

After establishing the molecular identity and functional significance of P57-activated MPA neurons, we next sought to identify their downstream projection circuits. Anterograde tracing with AAV2/9-EF1a-DIO-EYFP revealed that P57-activated MPA neurons project predominantly to the PVH, dorsomedial hypothalamus (DMH), and suprachiasmatic nucleus (SCH) (Fig. 4a, Fig. S7a-c). To determine which of these downstream projections mediating hypothermia, we combined retrograde AAV-Flpo virus with fDIO-Gq-mCherry viruses to selectively manipulate the MPA-PVH, MPA-DMH and MPA-SCH circuits. Chemogenetic activation of either MPA-PVH or MPA-DMH projections induced a rapid and significant drop in core body temperature, whereas activating the MPA-SCH projection had no measurable effect (Fig. S7d and e). Therefore, we further focused on the MPA-PVH and MPA-DMH pathways for further investigation.

**Fig. 4.**
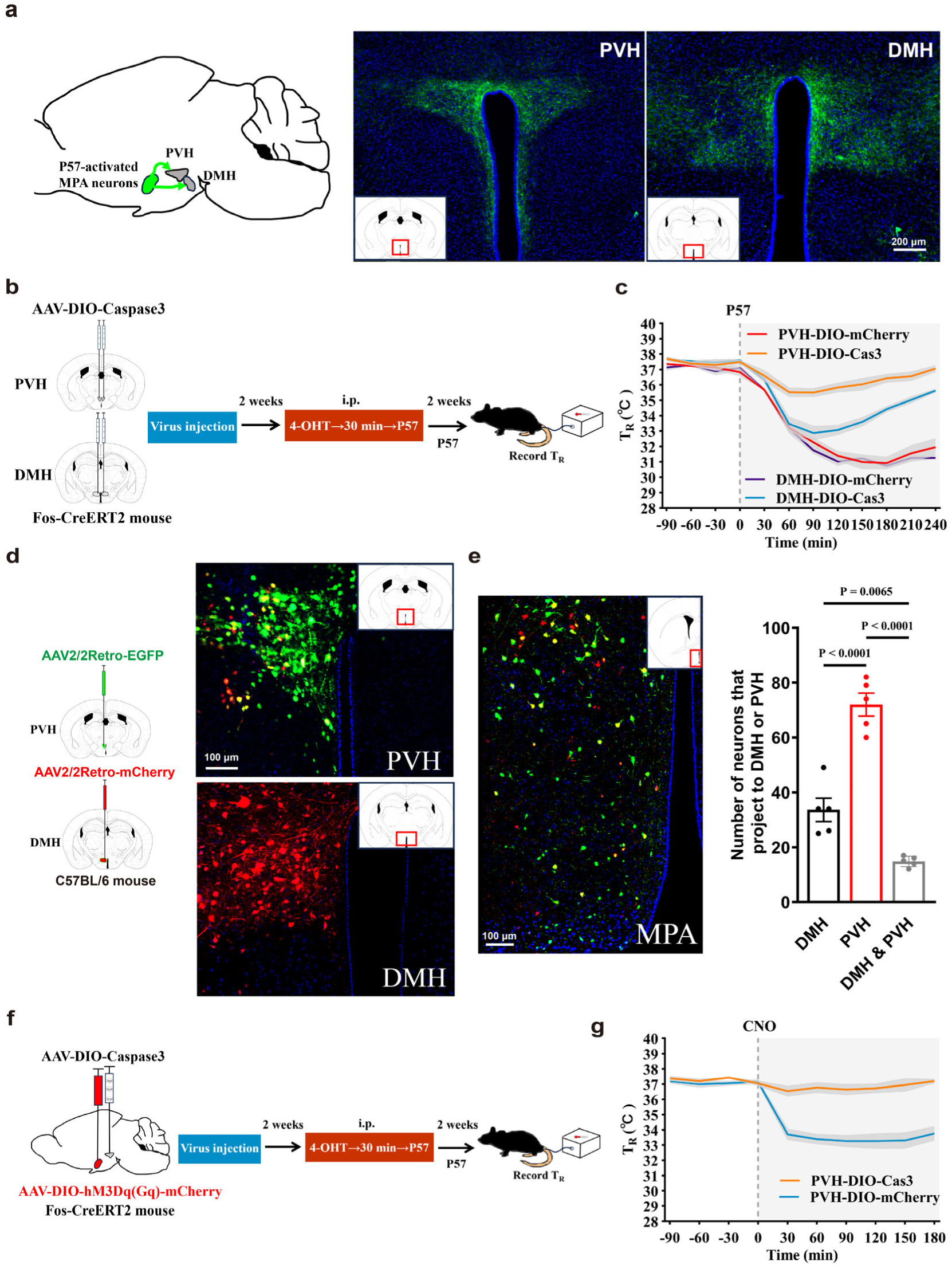
MPA-PVH circuit mediates P57-induced hypothermia. **a,** Representative images of projections to the PVH and DMH. **b,** Schematic shows the process of using AAV-DIO-Caspase3 to kill neurons in the PVH and DMH that respond to P57 in *Fos-CreERT2* mice. **c,** T_R_ of *Fos-CreERT2* mice before and after intraperitoneal injection of P57 after P57 responsive neurons killed (n=3 mice in each group). **d,** Schematic shows the process of using retroviruses to compare the projection of MPA to PVH and DMH (left), and the expression of the virus in PVH and DMH (right) (n=5 mice). **e,** Expression of the virus in **(d)** in MPA (left) with quantification (right). **f,** Schematic shows chemogenetic manipulation of neurons specifically activated by P57 in MPA while kill neurons in the PVH that respond to P57. **g,** T_R_ of *Fos-CreERT2* mice in **(f)** before and after intraperitoneal injection of CNO (n=4 mice in PVH-DIO-Cas3 group and 7 mice in PVH-DIO-mCherry group). Significant differences between groups were calculated using One-way ANOVA. Data are presented as mean values ± s.e.m.

To assess their functional relevance to P57-induced hypothermia, we expressed Cre-dependent Caspase3 in either PVH or DMH of *Fos-CreERT2* mice to selectively ablate neurons activated by P57 in these regions. Ablation of PVH neurons significantly blocked the P57-induced hypothermia, whereas loss of DMH neurons produced only a partial reduction in the hypothermic response (Fig. 4b and c, Fig. S7f and g). Consistent with these functional data, retrograde tracing revealed denser projections from MPA neurons to the PVH than to the DMH (Fig. 4d and e). Furthermore, combining PVH neuronal ablation with chemogenetic activation of P57-activated MPA neurons completely abolished the hypothermic effect (Fig. 4f and g). Collectively, these findings identify the MPA-PVH circuit as the primary pathway mediating P57-induced hypothermia.

### *Foxp2*^MPA^-targeted neuromodulation therapeutic hypothermia inhibits tumor growth

To determine whether hypothermia induced by *Foxp2*^MPA^ neurons is sufficient to confer therapeutic benefit and thus serve as a target for the neuromodulation-based hypothermia treatment, we evaluated its efficacy in disease models. Metabolic dysregulation is a defining feature of malignancy, wherein cancer cells reconfigure their biochemical pathways to orchestrate the bioenergetic and biosynthetic requirements for accelerated growth and sustained survival^40^. While prior studies have shown that cold exposure can suppress tumor growth, suggesting that tumor progression is temperature-sensitive, it remains unclear how ambient temperature versus its associated fluctuations in core body temperature independently contribute to these effects^5^. We therefore tested the therapeutic impact of *Foxp2*^MPA^ neuron induced hypothermia in tumor models and directly assessed the contribution of reduced body temperature per se to tumor progression. Following delivery of AAV-DIO-Gq-mCherry or control virus (AAV-DIO-mCherry) into the MPA of *Foxp2-Cre* mice, we allowed two weeks for viral expression before subcutaneous implantation of the MC38 murine colorectal tumor cells. Activating *Foxp2*^MPA^ neurons significantly slowed tumor progression, and the tumor averaged only 38.19 ± 6.55% volume and 34.00 ± 9.06% weight of those in the control group on day 14 (Fig. 5a). Twice-daily temperature monitoring showed that mice with activated *Foxp2*^MPA^ neurons remained continuously hypothermic (<34.5 °C) throughout most of the tumor-bearing period (Fig. 5b).

**Fig. 5.**
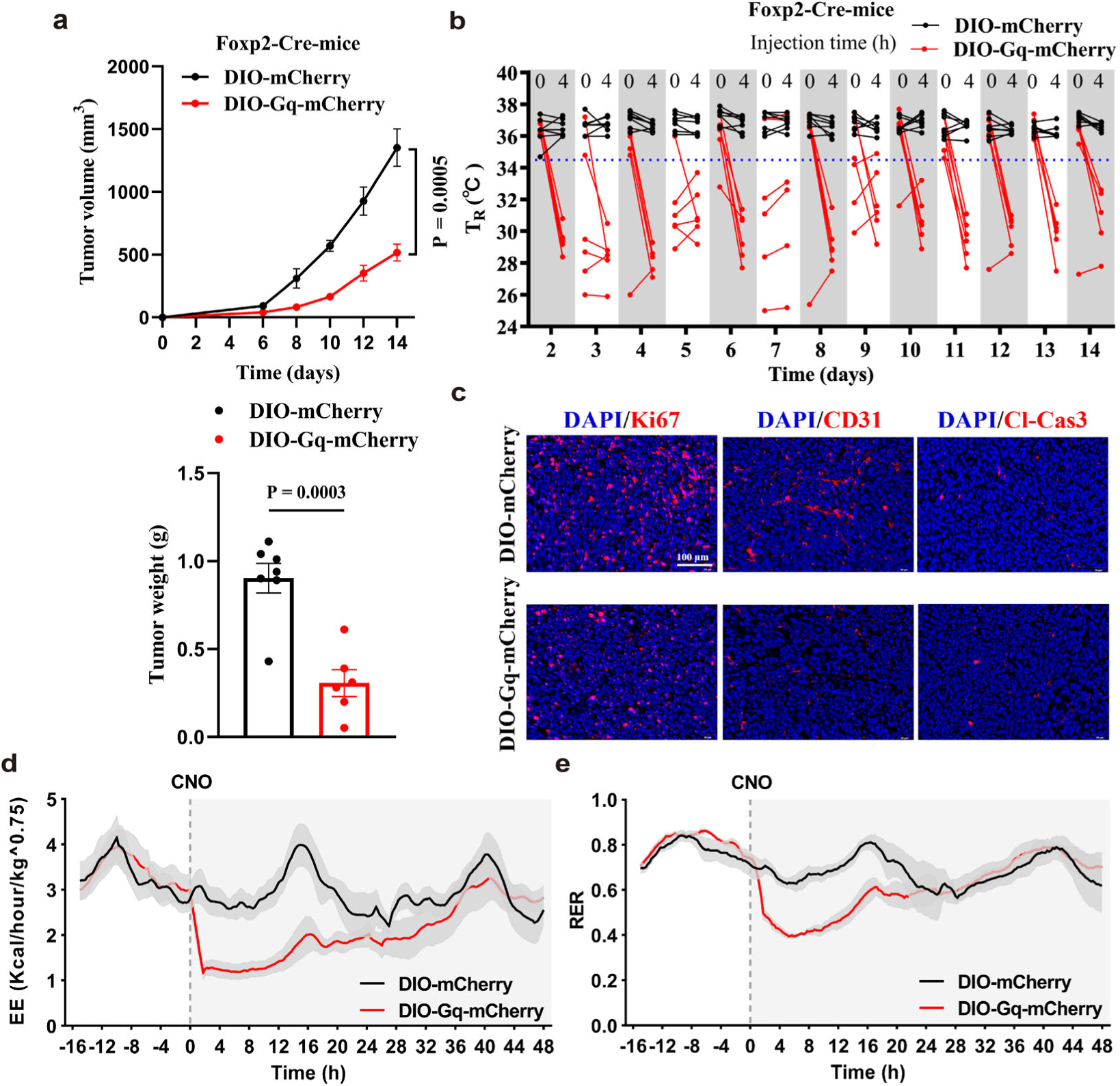
*Foxp2*^MPA^-targeted neuromodulation therapeutic hypothermia inhibits tumor growth. **a,** Tumor volume (top) and tumor weight (bottom, day 14) of *Foxp2-Cre* mice that received virus injection (n=6 mice in DIO-Gq-mCherry group and 7 mice in DIO-mCherry group). **b,** T_R_ of mice in **(a)** during experiment period. The administration and T_R_ measurement of mice are described in detail in the methods section. **c,** Immunofluorescence staining of tumor for Ki67, CD31 and Cleaved Caspase3 (Cl-Cas3). **d, e,** Energy expenditure **(d)** and respiratory exchange ratio **(e)** of *Foxp2-Cre* mice bearing tumor from day 7 to day 9 (n=4 mice in each group). The dashed line indicates the onset of CNO administration, shading indicates error bars. Student’s t test used was two-sided. Data are presented as mean values ± s.e.m.

Immunofluorescence staining of tumor tissues for classical markers of cell proliferation (Ki67), angiogenesis (CD31), and apoptosis (Cleaved-Caspase3, Cl-Cas3) revealed that hypothermic tumor-bearing mice displayed markedly reduced proliferation and lower microvessels area relative to controls, whereas no difference in cell apoptosis was detected between groups (Fig. 5c and Fig. S8a). These findings demonstrate that targeted activation of *Foxp2*^MPA^ neurons suppresses tumor progression primarily through inhibition of proliferation and angiogenesis, without directly affecting cancer cell apoptosis^5^.

To further assess the physiological impact of neuromodulation-based therapeutic hypothermia, we monitored metabolic dynamics in tumor-bearing mice (Day 7–Day 9) using metabolic cages during hypothermia induction. Activation of *Foxp2*^MPA^ neurons led to marked reductions in energy expenditure (EE), respiratory exchange ratio (RER), oxygen consumption, carbon dioxide production, water intake, and food intake (Fig. 5d and e, Fig. S8b-d). These coordinated changes indicate that the mice entered a global hypometabolic state.

We also asked whether global, non-selective activation of MPA neurons could reproduce this anti-tumor benefit in C57BL/6 mice (WT mice). Pan-neuronal excitation indeed slowed tumor growth, and by day 14, the tumor averaged 63.05 ± 17.30% volume (no significant difference) and 33.95 ± 12.05% weight of those in the control group (Fig. S9a and b) and lower core temperature (Fig. S9c). However, every animal developed marked polyuria (Fig. S9d), and two mice experienced fatal seizures. These adverse outcomes reinforce that global MPA activation produces uncontrollable side effects, underscoring the necessity for cell-type-specific targeting.

Collectively, these data confirm that neuronally induced hypothermia is sufficient to restrain tumor growth, indicating that *Foxp2*^MPA^ neurons induce hypothermia, imposing an energy bottleneck on rapidly proliferating cancer cells. Our findings validate *Foxp2*^MPA^ neurons as a viable neuromodulation target for therapeutic hypothermia. Encouragingly, analysis of spatial transcriptome datasets revealed regional distribution of these P57 specific activated neurons in MPA in human hypothalamus (Fig. S9e), which suggests that this population’s thermoregulatory role is evolutionarily conserved, providing a foundation for cross-species validation of the neuronal control of therapeutic hypothermia in tumor treatment.

## Discussion

Our study provides a real-time, whole-brain view of the hypothermic response induced by the small molecule P57, identifying the MPA as P57-specific activate region, and revealed the triggered neuronal population is characterized by the expression of *Foxp2* and *Prlr*. We further demonstrate that the newly identified *Foxp2*^MPA^ neurons are both sufficient and necessary for P57-induced hypothermia, and that the MPA-PVH circuit serves as the critical downstream conduit. Crucially, chemogenetic activation of *Foxp2*^MPA^ neurons establishes a proof-of-concept for neuromodulation-induced therapeutic hypothermia, providing direct evidence that reduced core temperature itself, separate from the systemic stress of ambient cold exposure, exerts therapeutic efficacy in oncology.

Building on previous work that defined the molecular targets of the *Hoodia gordonii*–derived compound P57^21^, we here extend those findings into a systems-level neurobiological framework. By mapping the neural circuits through which P57 induces hypothermia, we provide a template for neuromodulation strategies. A central implication is that *Foxp2*^MPA^ neurons represent a promising target for inducing therapeutic hypothermia. Because P57-induced hypothermia is a non-toxic, protective state, the *Foxp2*^MPA^ neurons are inherently suited for neuromodulation-based hypothermia treatment. Unlike traditional physical cooling, which triggers deleterious thermogenic defenses, targeting this population allows for a stable and controlled hypothermic state.

Notably, the hypothermic state induced by *Foxp2*^MPA^ neurons is not functionally or molecularly equivalent to natural hibernation or torpor. While we did not perform a systematic comparison, existing data distinguish *Foxp2*^MPA^ neurons from previously identified populations. Molecularly, our snRNA-seq dataset shows that *Foxp2*^MPA^ neurons are distributed across clusters distinct from *Adcyap1*^+^ neurons (linked to torpor-like states) or *Qrfp*^+^ neurons (linked to hibernation-like states)^12,15^. Phenotypically, *Foxp2*^MPA^ neuron activation maintains a minimum core body temperature of 29.7 ± 0.2 °C, which is warmer than the deep hibernation induced by *Qrfp*^+^ neurons (∼24 °C) but distinct from the torpor-like state induced by *Adcyap1*^+^ neurons (∼31 °C). These differences indicate that *Foxp2*^MPA^ neurons constitute a unique functional cluster optimized for therapeutic rather than extreme hypometabolism.

We further validated the potential of this *Foxp2*^MPA^-targeted strategy in an oncology model. Prolonged chemogenetic activation of these neurons maintained low body temperature, suppressed tumor proliferation and neovascularization without increasing apoptosis. Metabolic analysis indicates that the anti-tumor effect is driven by metabolic restraint rather than by cytotoxicity. However, a critical question is whether tumor suppression results directly from reduced core temperature or from pleiotropic physiological changes triggered by *Foxp2*^MPA^ neuron activation. As thermoregulatory neurons function within an integrated homeostatic network, their activation inevitably engages coordinated metabolic, endocrine, and vascular adaptations. Consequently, hypothermia cannot be fully decoupled from the systemic state it entails. While secondary adaptations cannot be formally excluded, we argue they are intrinsic to the hypothermic program rather than independent parallel effects. Given that temperature fundamentally dictates metabolic kinetics, we interpret this reduced core temperature as the primary driver of tumor inhibition. Thus, we propose that *Foxp2*^MPA^ neurons serve as a controllable node to induce a holistic hypometabolic state where hypothermia serves as the central mediator of anti-tumor efficacy.

In summary, we exploited P57 to identify *Foxp2*^MPA^ neurons as a key population for inducing therapeutic hypothermia and translated this into a neuromodulation-based therapeutic hypothermia that suppresses tumor growth. By demonstrating that controlled hypothermia alone is sufficient to restrain cancer, we broaden the indications for hypothermia-based interventions. Future efforts should systematically investigate the optimal depth and duration of hypothermia across diverse cancer types, including both solid and hematological malignancies, while expanding beyond subcutaneous models to evaluate effects on spontaneous and metastatic disease. Furthermore, it will be critical to evaluate the long-term systemic impact of sustained hypothermia on factors such as metabolic homeostasis, immune function, and cognitive performance. Integrating closed-loop brain machine interfaces to replace viral-based delivery, will also be essential to enable non-invasive, on-demand precision hypothermia. Addressing these limitations will be critical for advancing toward the clinical translation of neuronally induced therapeutic hypothermia. More broadly, the principle of re-tuning neural activity to reshape physiological states may extend neuromodulation beyond classical neurological disorders, offering drug-free therapeutic options for chronic diseases rooted in homeostatic imbalance.

## Methods

### Cell culture

The HEK293T cell line (ATCC, Virginia, USA) was cultured with DMEM (Gibco) containing 10% fetal bovine serum (FBS, Biosera) and 1% penicillin and streptomycin. MC38 cells were from Professor Jun Liu’s Lab in Johns Hopkins University as a gift. MC38 cells were cultured in RIPM-1640 (Gibco) containing 10% FBS. All cells were maintained in 5% CO_2_ at 37 °C.

### Mouse studies

All mouse care and experimentation were ethically performed according to procedures approved by the Institutional Animal Care and Use Committee at Fudan University. Adult (6-10-week-old) male C57BL/6 mice were purchased from Shanghai JieSiJie Laboratory Animal Company. For whole brain imaging and virus injection, we used adult (6-10-week-old) male and female *Fos-CreERT2;tdTomato* mice (Jackson Lab), *Fos-CreERT2* mice (Jackson Lab), and *Foxp2-Cre* mice (Jackson Lab). All mice were caged at 22 ± 2 °C under 12 h–12 h light–dark cycles.

No statistical methods were used to predetermine the sample size. Mice were randomly assigned to experimental groups before surgery.

### Chemicals and Drug preparation

P57^41^ was provided by Xiaheng Zhang’s group by total synthesis. 4-Hydroxytamoxifen (4-OHT) was purchased from TargetMol (T6743). Clozapine N-oxide (CNO) was purchased from AbMole (M7684). Pyridoxal (PL) was purchased from J&K Scientific (322481).

For intraperitoneal injection (i.p.), P57 solution (25 mg/kg for mice) was prepared by initially dissolving 2.5 mg P57 in 10 μL DMSO, and then adding 990 μL 10% castor oil (1:9 mixture of castor oil: PBS). 4-OHT solution (50 mg/kg for mice) was prepared by initially dissolving in 500 μL 100% ethanol and 450 μL mixture oil (1:4 mixture of castor oil: sunflower oil), then removing the ethanol via vacuum centrifugation and adding the mixture oil to a final concentration of 6.25 mg/mL. CNO solution (3 mg/kg for mice) was prepared by initially dissolving 0.3 mg CNO in 10 μL DMSO, and then adding 990 μL isotonic saline. PL solution (300 mg/kg for mice) was prepared by dissolving 30 mg PL in 1 mL PBS. Vehicle control means a mixture of 1% DMSO, 10% castor oil, and 89% PBS for P57, or PBS for PL.

For cell experiments, PL at the corresponding concentration was dissolved in PBS. Vehicle control means PBS.

### Virus injection

AAV2/2-hSyn-DIO-hM3Dq(Gq)-mCherry-WPRE-pA (Taitool, S0192-2), AAV2/9-hEF1a-DIO-mCherry-P2A-TetTox-WPRE-pA (Taitool, S0506-9-H5), AAV2/9-hEF1a-DIO-EYFP-WPRE-pA (BrainVTA, PT-0012), scAAV2/2Retro-hSyn-Flpo-pA (Taitool, S0294-2R), AAV2/9-hEF1a-fDIO-hM3Dq(Gq)-mCherry-ER2-WPRE-pA (Taitool, S0337-9), AAV2/9-hSyn-DIO-mCherry-WPRE-pA (Taitool, S0240-9), AAV2/9-CAG-DIO-taCaspase3-TEVp-WPRE-pA (Taitool, S0236-9), AAV2/2Retro-CMV_bGI-EGFP-WPRE-pA (Taitool, S0263-2RP), AAV2/2Retro-CMV_bGI-mCherry-WPRE-pA (Taitool, S0244-2R), rAAV-hSyn-hM3D(Gq)-EGFP-WPRE-pA (BrainVTA, PT-0152) and AAV2/9-hSyn-EGFP-WPRE-pA (Taitool, S0237-9-H20) were used for viruses injection. All viruses were diluted with PBS to a final concentration of 5×10^12^ per mL before stereotaxic delivery into the mouse brain.

For injection of AAV2/2-DIO-hM3Dq(Gq)-mCherry, AAV2/9-DIO-mCherry, AAV2/9-DIO-TetTox, or AAV-fDIO-hM3Dq(Gq)-mCherry into MPA, *Fos-CreERT2* mice or *Foxp2-Cre* mice were anaesthetized with isoflurane and positioned in a stereotaxic frame (RWD Life Science). 480 nL (120 nL per site) of virus was delivered into MPA (AP, 0.2 mm; ML, ±0.25 mm; DV, -5.4/-5.6 mm) at a controlled rate of 2 nL per second using a Hamilton needle syringe. For AAV2/9-hEF1a-DIO-EYFP, only one side of the MPA (AP, 0.2 mm; ML, -0.25 mm; DV, -5.4/-5.6 mm) was injected with the virus. For AAV2/2Retro- EGFP and AAV2/2Retro-mCherry, only one side of the PVH (AP, -0.95 mm; ML, -0.2 mm; DV, -4.8 mm) or DMH (AP, -1.7 mm; ML, -0.3 mm; DV, -5.0 mm) was injected with the virus. The needle was kept in place for 10 min after injection. The waiting period for recovery and virus expression for the experiments was 2 weeks.

For the injection of scAAV2/2Retro-Flpo, AAV-DIO-Caspase3 or AAV2/9-DIO-mCherry, *Fos-CreERT2* mice or C57BL/6 mice were anaesthetized with isoflurane and positioned in a stereotaxic frame. 120 nL of virus per site was delivered into PVH (AP, -0.95 mm; ML, ± 0.2 mm; DV, -4.8 mm), DMH (AP, -1.7 mm; ML, ± 0.3 mm; DV, -5.0 mm) and SCH (AP, -0.7 mm; ML, ± 0.1 mm; DV, -5.6 mm) at a controlled rate of 2 nL per second using a Hamilton needle syringe.

### Mouse body temperature recording

The mouse rectal temperature (T_R_) was recorded by an animal temperature measuring apparatus (FT3400, Kew Basis) every 30 minutes. All mice were adapted for 5 days before the test. All animal experiments were conducted at ambient temperature at 20∼24°C.

### Functional ultrasound (FUS) imaging

#### Mice craniotomy

A headpost with an imaging window was surgically implanted in C57BL/6 mice for head fixation, as described in detail previously^42^. Briefly, Mice were anesthetized with isoflurane, their hair was shaved off, and the skin was disinfected with a cutoff. Lidocaine was given to the scalp and then the scalp was cut open. A metal headframe was fixed to the skull with dental cement (Super-bond C&B, Nissin). Carefully grind the skull with a drill bit until it can be easily removed with tweezers while keeping the dura mater intact. The imaging window of the metal headframe was covered with Polymethylpentene (Merck, GF42802129). The skull, headframe, and Polymethylpentene were wrapped with photocuring resin to form a seal, and artificial cerebrospinal fluid (ACSF, containing 127 mM NaCl, 2.5 mM KCl, 25 mM NaHCO_3_, 1.25 mM NaH_2_PO_4_, 2.0 mM CaCl_2_, 1.0 mM MgCl_2_, and 25 mM Glucose) was injected.

#### Habituation, imaging, and data analysis

FUS acquisitions were collected using an Iconeus One functional ultrasound imaging system (Iconeus One, Iconeus, Paris, France) tailored for animal studies. Seven days after the surgical recovery, the heads of the awake mice were fixed on a fixed bracket, allowing their limbs to move freely. The Iconeus One functional ultrasound imaging system was run using the same process as in the formal experiment, the mice were trained for 5 days to adapt. During the formal experiment, the cerebral blood flow activity data of mice for 10 minutes were first recorded as the baseline. Then, P57 or vehicle was intraperitoneally injected into the mice, and the signals were collected for another 40 minutes.

A 4D ultrasonic probe (IcoPrime-4D MultiArray 15 MHz, Iconeus, Paris, France) connected to an ultrafast ultrasound scanner (Iconeus One, 256 channels, Iconeus, Paris, France) was used. This probe consists of four linear arrays, each containing 64 elements, operating at a center frequency of 15 MHz with a 110 μm pitch. It provides a field of view of 7 mm in width × 8 mm in length, and up to 20 mm in depth, with a plane thickness of ∼500 μm and an in-plane spatial resolution of 100 μm × 100 μm. Throughout the experiments, the probe was securely mounted on a 4-axis motorized stage, providing movement with four degrees-of-freedom (3-dimensional translation and rotation along the vertical axis). This setup ensured precise positioning and orientation, facilitating accurate alignment and co-registration of each mouse’s 3D whole-brain image with the Allen Mouse Common Coordinate Framework using a fully automated registration mode. With its simultaneous multi-slicing technology, the entire mouse brain was covered in 2.4 seconds. The skull of the mouse was covered with isotonic coupling gel. Four power Doppler (PD) images (one per array) were simultaneously obtained from 4 × 200 compounded frames acquired at 500 Hz using 4 × 12 tilted plane waves acquired at a pulse repetition frequency of 4 kHz (-12°, -9.82°, -7.64°, -5.46°, -3.28°, -1.1°, 1.1°, 3.28°, 5.46°, 7.64°, 9.82°, 12°). To compensate for the limited lateral aperture size (64 elements compared to 128 in conventional linear arrays), we used a trapezoidal beamforming grid with *θ_max_* =12°, allowing the field of view to be extended on both sides and enabling the retrieval of deeper lateral brain regions. Each block of 0.4 s was filtered using a Singular Value Decomposition (SVD) clutter filter to separate tissue signal from blood signal and form a PD image^43^. The dedicated imaging sequence and the live Doppler reconstruction procedure were implemented in a live acquisition software (IcoScan, Iconeus, Paris, France).

The registration and cerebral blood volume (CBV) signals analysis of the mouse brain regions was all completed in the Icostudio 2.2 software and exported^22^. Taking the CBV data 1 to 9 minutes before intraperitoneal injection as the average baseline, the CBV at each time point was divided by the average baseline to obtain the relative CBV (rCBV), and the outliers were eliminated using the interquartile range method. Then calculate ΔrCBV according to the following formula: ΔrCBV = (average(rCBV_P57)-average(rCBV_Vehicle))/average(rCBV_Vehicle), average(rCBV_P57) refers to the average value of rCBV in each mouse at each time point in P57 group and average(rCBV_Vehicle) refers to the average value of rCBV in each mouse at each time point in Vehicle group. Data processing and visualization are implemented using Python 3.13.0. 3D rendering of each region of the hypothalamus was performed using brainrender via brainglobe-heatmap^44^.

#### Brain processing, immunofluorescence, and imaging

Mice were deeply anesthetized with isoflurane, and perfused transcardially with pre-cooled PBS and 4% paraformaldehyde (PFA). Brains were fixed in 4% PFA at 4℃ for 24 hours and then sliced into 50 μm coronal sections using a vibratome (Leica). Brain sections were washed three times with PBS and treated in 0.2% Triton X-100 in PBS for 1 h at room temperature (RT), then blocked with 0.05% Triton X-100, 10% bovine serum albumin (BSA) in PBS for 1 h at RT. Brain sections were then incubated with primary antibody Rabbit anti-Foxp2 (1:200, Abcam, ab307505), Mouse anti-Prlr (1:500, proteintech, 67292-1-Ig), Guinea pig anti-cFos (1:5000, Synaptic systems, 226308), Rabbit anti-Pdxk (1:1000, proteintech, 15309-1-AP), Rabbit anti-Acadsb (1:200, proteintech, 13122-1-AP), Rabbit anti-NeuN (1:1000, proteintech, 26975-1-AP) or Mouse anti-pERK1/2 (T202/Y204) (1:1000, CST, 9106) in PBS with 0.05% Triton X-100 for 24∼48 hours at 4℃. Tissues were rinsed in PBS for three times, and incubated with secondary antibody solution, anti-Guinea pig 488 (1:1000, Thermofisher, A11073), anti-Rabbit 488 (1:1000, Thermofisher, A11034), anti-rabbit 594 (1:1000, Thermofisher, A11012), anti-mouse 647 (1:1000, Thermofisher, A21236) or anti-rabbit 647 (1:1000, Thermofisher, A21245) in PBS for 2 hours at RT, then rinsed with PBS for three times and mounted onto slides, dried, and covered under DAPI Fluoromount-G^®^ (SoutherBiotech, 0100-20). Sections were imaged with an Olympus VS120 slide scanning microscope or PanoBrain slide scanning system (Meca Scientific). Confocal images were acquired with an Olympus FV3000 microscope. Images were analyzed in ImageJ (FIJI) and QuPath.

#### Labeling neurons

*Fos-CreERT2;tdTomato* mice and *Fos-CreERT2* mice were used to label neurons specifically activated by P57. These mice were intraperitoneally injected with isotonic saline at the same time every day for five days before receiving the intraperitoneal injection of the P57 to adapt to the injection stimulation. On the day of labeling, 4-OHT was injected 30 minutes in advance, followed by P57 or vehicle. *Fos* and *CreERT2* were expressed in the neurons activated by P57. In the presence of 4-OHT, CreERT2 translocates into the nucleus, enabling *tdTomato* or other Cre-dependent AAV-DIO viruses to be specifically expressed in *FOS*-expressing cells activated by P57.

#### Image registration to the Allen Brain Atlas and analysis

The preparation of *Fos-CreERT2;tdTomato* mice brain slices was as described above. For the methodology, whole-slide images of brain tissue sections were first acquired using the PanoBrain slide scanning system. All subsequent image processing and analysis were conducted within the Panolyzer software environment. The analysis pipeline consisted of four main steps: (1) Initial registration of the histological images to the Allen Mouse Brain Atlas^45^ was performed using the DeepSlice method^46^. (2) To achieve a more precise anatomical alignment, a non-linear b-spline registration algorithm was applied for fine refinement^47^. (3) Following registration, TRAPed cells were initially quantified on the aligned images by applying a threshold-based segmentation algorithm. (4) The results from the automated cell quantification were then manually reviewed and corrected to yield the final cell statistics. The registration and statistics of other mouse brain slices were conducted using ImageJ (FIJI) and QuPath.

#### Mouse hypothalamus snRNA-seq

Mice were deeply anesthetized with isoflurane and perfused transcardially with pre-cooled PBS. The brain was removed, and the complete hypothalamus was isolated. It was directly quick-frozen in liquid nitrogen for preservation and used in snRNA-seq. Single nuclei were captured and barcoded for whole-transcriptome libraries using the Singleron platform according to the manufacturer’s recommendations, collecting one library of approximately 10,000 nuclei from each of the 3 mice. Single-cell suspensions (2×10^5^ cells/mL in PBS, HyClone) were loaded onto a microwell chip using the Singleron Matrix NEO® Single Cell Processing System. Barcoding Beads were collected from the chip and used for reverse transcription (RT) to obtain cDNA, followed by PCR amplification. The amplified cDNA was subsequently fragmented and ligated with sequencing adapters. Libraries were constructed according to the protocol of the GEXSCOPE® Single Cell RNA Library Kits (Singleron)^48^. Individual libraries were diluted to 4 nM, pooled, and sequenced on Illumina Novaseq 6000 with 150-bp paired-end reads.

Raw sequencing reads were processed to generate the gene expression matrix using the CeleScope pipeline (v2.0.7) https://github.com/singleron-RD/CeleScope). Initial quality control and UMI extraction followed the standard CeleScope workflow. Briefly, raw reads were first processed using Cutadapt (v1.17) to remove low-quality reads, poly-A tails, and adapter sequences. Cell barcodes and Unique Molecular Identifiers (UMI) were subsequently extracted. Given that the mouse samples expressed the exogenous hM3Dq receptor gene, the standard mouse reference genome was customized to ensure comprehensive mapping. Following alignment, UMI counts and gene counts for every single cell were acquired using the featureCounts (v2.0.1) software. The resulting expression matrix files were used for all subsequent quality filtering and downstream bioinformatic analyses.

#### Data Analysis, Quality Control, Dimension Reduction, and Clustering

Data analysis was performed using R software (v4.3.1) and the Seurat package (v5.4.0). The initial step involved quality control of the raw UMI count matrix. Cells and genes were filtered based on specific criteria: only genes detected in more than three cells were retained, and cells were kept if they had gene counts exceeding 200 but below 7,500, UMI counts greater than 1,000 but less than 60,000, and a mitochondrial gene content lower than 15%. This filtration process resulted in a final dataset comprising 26,087 cells and 24,275 genes for subsequent analysis. The UMI count matrix was normalized and scaled using the NormalizeData and ScaleData functions, respectively. Dimensionality reduction was initiated by identifying the top 3,000 highly variable genes using the FindVariableFeatures function, which were then used as input for principal component analysis (PCA) via the RunPCA function. The optimal number of principal components (PCs) for downstream analysis, determined to be 15, was identified by evaluating the PCA results with the ElbowPlot and DimHeatmap functions. Cell clustering was carried out using the top 15 principal components. The FindNeighbors and FindClusters functions (resolution = 0.5) were employed to group the cells into distinct clusters. Finally, the cell clusters were visualized in a two-dimensional space using the Uniform Manifold Approximation and Projection (UMAP) algorithm.

#### Cell Type Annotation

The marker genes of each cluster were identified using the FindAllMarkers function. The expression patterns of the definitive marker genes used to assign cell type identities to each cluster were visualized using the DotPlot function from Seurat (v5.3.0). Marker genes for each cell type were listed here: Slc17a6 for Glutamatergic neurons, Gad2 for GABA neurons, Mbp for Oligodendrocyte cells, Vtn for Mural cells, Aldoc for Astrocyte cells, Cx3cr1 for Microglial cells, Ccdc153 for Ependymal cells, Pdgfra for OPC cells, Flt1 for Endothelial cells, Rax for Tanycyte, Dcn for Fibroblast.

#### Mouse tumor models

*Foxp2-Cre* mice were inoculated subcutaneously on the flank with 5 × 10^5^ MC38 colorectal cancer cells after 2 weeks of AAV-DIO-hM3Dq(Gq)-mCherry or AAV-DIO-mCherry injection in MPA. C57BL/6 mice were inoculated subcutaneously at flank with 5 × 10^5^ MC38 colorectal cancer cells after 2 weeks of AAV-hM3Dq(Gq)-mCherry or AAV-mCherry injection in MPA.

The day of tumor cell implantation in mice was recorded as Day 0. For intraperitoneal injection of CNO (3 mg/kg), starting from Day 2, the rectal temperature of mice was measured before administration each day and recorded as the T_R_ at hour 0. For the DIO-mCherry group and the mCherry group of mice, CNO was administered daily. For the mice in the DIO-Gq-mCherry group and Gq-EGFP group, CNO was given when their T_R_ was higher than 34.5℃ at 0 h; otherwise, it was not given. The T_R_ of the mice was measured again 4 hours after administration. All mice were stopped from being administered the CNO for one day on Day 7. Starting from Day 6, the tumor volume was calculated as follows: 1/2 × length × width^2^ every two days. On day 14, 4 hours after CNO administration, the tissues of the mice were taken for subsequent experiments, and the tumor weights were weighed.

#### Metabolic Studies

Tumor-bearing *Foxp2-Cre* mice were maintained individually in a metabolism chamber (Panlab Oxylet System, Harvard Apparatus) with free access to food and water for 72 hours. Mice were housed for 24 hours for adaptation. On the second day, mice were intraperitoneally injected with CNO. Metabolic parameters, including O_2_ consumption (VO_2_), CO_2_ production (VCO_2_), respiratory exchange ratio (RER), energy expenditure, food intake, and water intake, were recorded at 15 min intervals in a standard light-dark cycle. The respiratory quotient is the ratio of carbon dioxide production to oxygen consumption, RER = VCO_2_/VO_2_. Energy expenditure (EE) was calculated as follows: EE = 3.815*VO_2_ + 1.232*VCO_2_ according to the manufacturer’s instruction manual. Data analysis was conducted using METABOLISM v3.0.01 software.

#### Immunofluorescence

For immunofluorescence, tumor tissues were fixed by 4% PFA and then were dehydrated, embedded in paraffin, and cut into serial sections at 5 μm. Paraffin sections were dewaxed with xylene and rehydrated with gradient ethanol, then continuously heated in Tris-EDTA for 15 minutes for antigen remediation. After blocking with 3% BSA in PBS for 1 h at RT, sections were then incubated with primary antibody Rabbit anti-Ki67 (1:1000, Proteintech, 28074-1-AP), Mouse anti-CD31 (1:50, Biosciences, 550389) or Rabbit anti-cleaved caspase3 (1:1000, CST, 9664T) in PBS with 1% BSA for 24 hours at 4℃. Tissues were rinsed in PBS for three times, and incubated with secondary antibody solution, anti-rabbit 594 (1:1000, Thermofisher, A11012), anti-mouse 594 (1:1000, Thermofisher, A11032) in PBS for 2 hours at RT, then rinsed with PBS for three times, dried, and covered with DAPI Fluoromount-G^®^ (SoutherBiotech, 0100-20). Sections were imaged with an Olympus VS120 slide scanning microscope.

#### Western immunoblotting

HEK293T cells were lysed in a buffer containing 20 mM Tris-HCl (pH=7.4), 100 mM KCl, 1% Triton X-100, and a protease inhibitor cocktail. The following antibodies were used for immunoblotting: ACADSB (1:1000, Proteintech, 13122-1-AP), HSP90 (1:5000, Proteintech, 60318-1-Ig). Nitrocellulose membranes were incubated overnight with primary antibody at 4°C, followed by three times washes in PBST (PBS with 0.1% Triton X-100, vol/vol) before incubation in secondary antibodies. Peroxidase AffiniPure Goat Anti-Rabbit IgG (H+L) (1:10000, Jackson ImmunoResearch Inc, 111-035-003) and Goat Anti-Mouse IgG (H+L) (1:10000, Jackson ImmunoResearch Inc, 115-005-003) for 1 hour at room temperature. Images were captured using the ChemiScope system (Clinx Science Instruments Co. Ltd).

### PELSA (Peptide-centric Local Stability Assay)

#### Tissue collection and homogenization

As described above, obtain the hypothalamic tissue of mice. For lysis, frozen hypothalamic tissues were homogenized in PBS supplemented with 1% (v/v) protease inhibitor cocktail using a bead-beater homogenizer with a mixture of 2 mm and 3 mm stainless-steel beads. Homogenization was performed in three cycles of 30 s at 5 m/s, with a 30 s pause between cycles. The homogenate was centrifuged at 20,000 × g for 10 min at 4 °C to remove debris, and the protein concentration in the supernatant was determined using the Pierce™ 660 nm Protein Assay Kit (Thermo Fisher Scientific). The lysate was adjusted to a final protein concentration of 1 mg/mL.

#### Limited proteolysis and peptide preparation

Fifty microliters of lysate (50 μg protein) were incubated with 1 mM PL (treatment) or an equal volume of vehicle (control) at 25 °C for 30 min. Limited proteolysis was then initiated by adding 25 μg trypsin (Sigma-Aldrich), resulting in an enzyme-to-substrate ratio of 1:2 (w/w). Digestion was allowed to proceed for exactly 1 min at 37 °C and was quenched by adding a three-fold volume of 8 M guanidine hydrochloride (GdmCl). Samples were reduced with 10 mM tris(2-carboxyethyl) phosphine (TCEP) and alkylated with 40 mM chloroacetamide (CAA) at 95 °C for 5 min. To remove undigested proteins and large fragments, the mixture was centrifuged through a 10 kDa molecular-weight cutoff filter at 14,000 × g for 1 h. The filtrate was collected, and the filter was washed with 200 μL of 60 mM HEPES (pH 8.0) followed by another centrifugation step. The combined filtrates were acidified with 1% trifluoroacetic acid (TFA).

#### Peptide desalting and LC-MS/MS analysis

Acidified peptides were desalted using C18 StageTips prepared in-house with Sep-Pak® C18 material (Waters). After washing with 0.1% TFA, peptides were eluted with 80% acetonitrile/0.1% TFA, dried in a SpeedVac concentrator (Thermo Fisher Scientific), and reconstituted in 0.1% formic acid.

DIA analysis was performed on an Orbitrap Exploris 480 mass spectrometer coupled online to a Dionex Ultimate 3000 RSLC micro-system for microflow liquid chromatography. Peptides were separated by microflow liquid chromatography using a 15 cm × 1 mm i.d. reversed-phase column (ACQUITY UPLC Peptide CSH C18, 130 Å, 1.7 μm; Waters). A total of 10 μg of peptides were loaded onto the column and eluted at a flow rate of 50 μL/min with a binary solvent system: solvent A (0.1% formic acid in water) and solvent B (0.1% formic acid in 80% acetonitrile). The separation gradient was as follows: 6% to 32% B over 80 min, followed by an increase to 45% B over 14 min.

Ion mobility separation was performed using a high-field asymmetric waveform ion mobility spectrometry (FAIMS) interface. Analyses were carried out using a single compensation voltage (CV) of –45 V, with the FAIMS carrier gas (N2) maintained at a flow rate of 3.5 L/min and a temperature of 100 °C.

Mass spectrometry data were acquired in DIA mode. Full MS1 scans were acquired at a resolution of 120,000 (at m/z 200) over the m/z range of 350–1400, with an automatic gain control (AGC) target of 3 × 10^6^ and a maximum injection time of 45 ms. For DIA scans, the mass range m/z 400–1000 was divided into 24 variable-width windows. MS2 scans were acquired at a resolution of 30,000 (at m/z 200) with an AGC target of 2 × 10^6^, using automatic maximum injection time and a normalized higher-energy collisional dissociation (HCD) collision energy of 30%. The first fixed mass was set to m/z 300.

#### CETSA (Cell thermal shift assay)

The HEK293T cell pellet was resuspended in PBS containing protease inhibitors and then repeatedly frozen and thawed in liquid nitrogen for four times to break the cells. Centrifuge 20,000 g at 4 ℃ for 10 minutes and collect the supernatant as the protein solution. The protein solution was evenly divided and incubated with the PL (final concentration at 1 mM in PBS) or PBS at room temperature for 1 hour. Then, each solution was incubated at the set temperatures (43 ℃, 46 ℃, 49 ℃, 52 ℃, 55 ℃, 58 ℃) for 3 minutes. After that, they were taken out and placed in an ice box to stand for 3 minutes, then returned to room temperature. Then, centrifuge at 20,000 g at 4 ℃ for 10 minutes, carefully aspirate the supernatant, add it to the Loading buffer and boil the sample for 10 minutes at 98 ℃ for Western immunoblotting.

#### Human hypothalamic spatial transcriptome data

Spatial transcriptomics data were obtained from the study “A comprehensive spatio-cellular map of the human hypothalamus” (GSE278848). The original experiments were performed on formalin-fixed, paraffin-embedded (FFPE) human hypothalamus sections (5 μm thickness) using the 10x Genomics Visium CytAssist v.2 workflow. The core data processing, including demultiplexing and read alignment, was performed using SpaceRanger v.2.0.0 (10x Genomics). Reads were aligned to the Visium human transcriptome probe set v.2 (reference: GRCh38 2020-A). This pipeline mapped unique molecular identifiers (UMIs) to the Visium spot coordinates, generating raw count matrices and spatial barcode coordinates for each section.

Quality control metrics from the original study indicated high data quality, with a median of 7,105 counts and 3,560 detected genes per spot across nine samples. Prior to analysis, the raw count matrices and spatial barcode coordinates for each sample were inspected. Spots previously identified as artifacts by the original authors (e.g., through Loupe Browser inspection due to aberrant counts) were excluded to ensure data integrity.

For gene expression visualization and spatial distribution analysis, the filtered count matrices were imported into R (v.4.5.0) using the Seurat package (v.5.2.0). The data were then log-normalized using the standard Seurat workflow to account for variations in sequencing depth among spots. The spatial expression patterns of target genes, were visualized by mapping normalized expression values onto the corresponding Visium spot coordinates of the tissue section. SpatialFeaturePlotfunction was used to generate high-resolution spatial maps of gene expression across the hypothalamic tissue.

## Data Availability

Data supporting the findings of this study are available from the corresponding authors upon reasonable.

## Author contributions

Y.D., W.J. and L.X. conceived and directed the project. G.B. and L.Z. did experiments and analyzed and prepared data. G.B., J.C. and J.C. contributed to neurobiological experiments. G.B., J.C., Z.Z., X.L., Y.Z. and G.L. contributed to tumor model. L.Z., G.B. and M.L. analyzed the snRNA sequencing data. M.Y., T.Y. and G.B. contributed to identification of target protein. D.M. and Z.Z advised on the study. B.Y., L.W. and X.Z. contributed to chemical synthesis. The manuscript was written by G.B., L.Z., L.X., W.J. and Y.D.

## Supplementary Figure legends

**Fig. S1.**
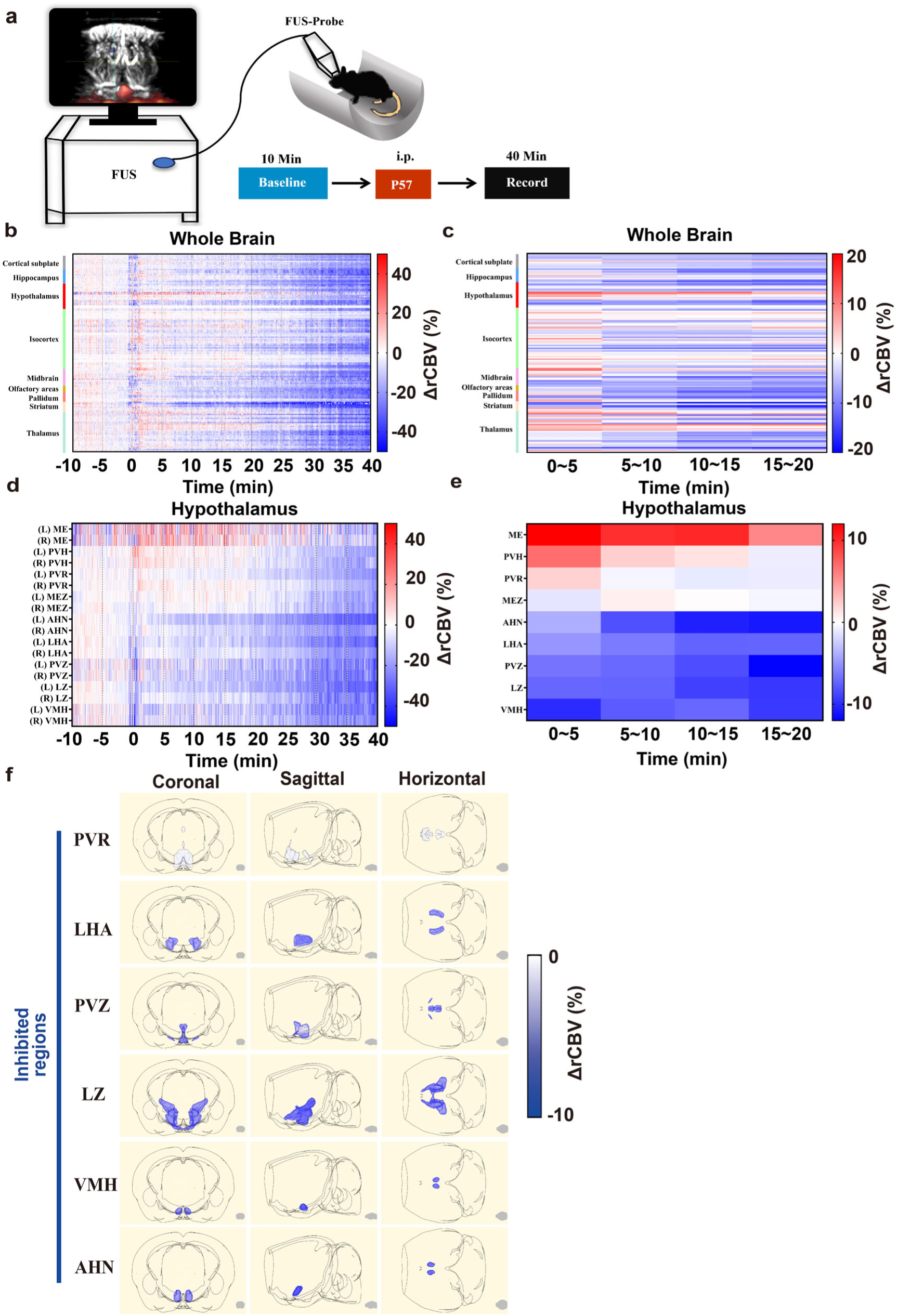
Changes in cerebral blood volume in various brain regions of mice after administration of P57. **a,** Schematic of the Functional ultrasound (FUS) experiment design for awake C57BL/6 mice (n = 4 mice for P57 group and 5 mice for Vehicle group). **b,** Real-time ΔrCBV of brain throughout the experiment. **c,** ΔrCBV at different time periods within 20 minutes after administration. **d,** Real-time ΔrCBV of hypothalamus throughout the experiment. **e,** ΔrCBV of hypothalamus at different time periods within 20 minutes of administration**. f,** Different views of the 3D rendering results of various P57-inhibited brain regions in the hypothalamus within 5 to 10 minutes after administration. Periventricular region (PVR), Lateral hypothalamic area (LHA), Periventricular zone (PVZ), Hypothalamic lateral zone (LZ), Ventromedial hypothalamic nucleus (VMH), Anterior hypothalamic nucleus (AHN).

**Fig. S2.**
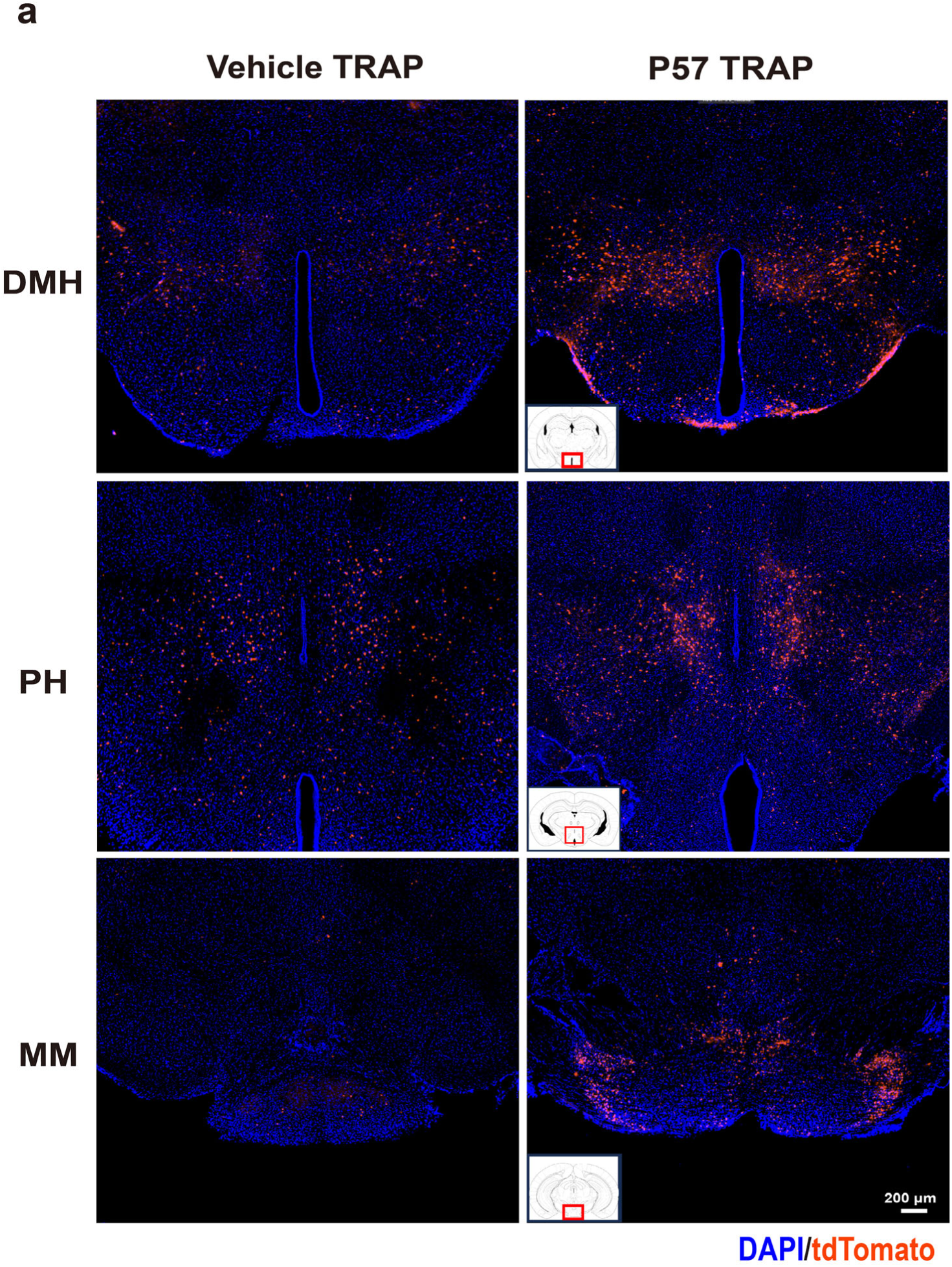
The tdTomato expression in PH and DMH. Representative slice of posterior hypothalamic nucleus (PH), dorsomedial nucleus (DMH) and mammillary body (MM) in P57 group and Vehicle group in *Fos-CreERT2;tdTomato* mice.

**Fig. S3.**
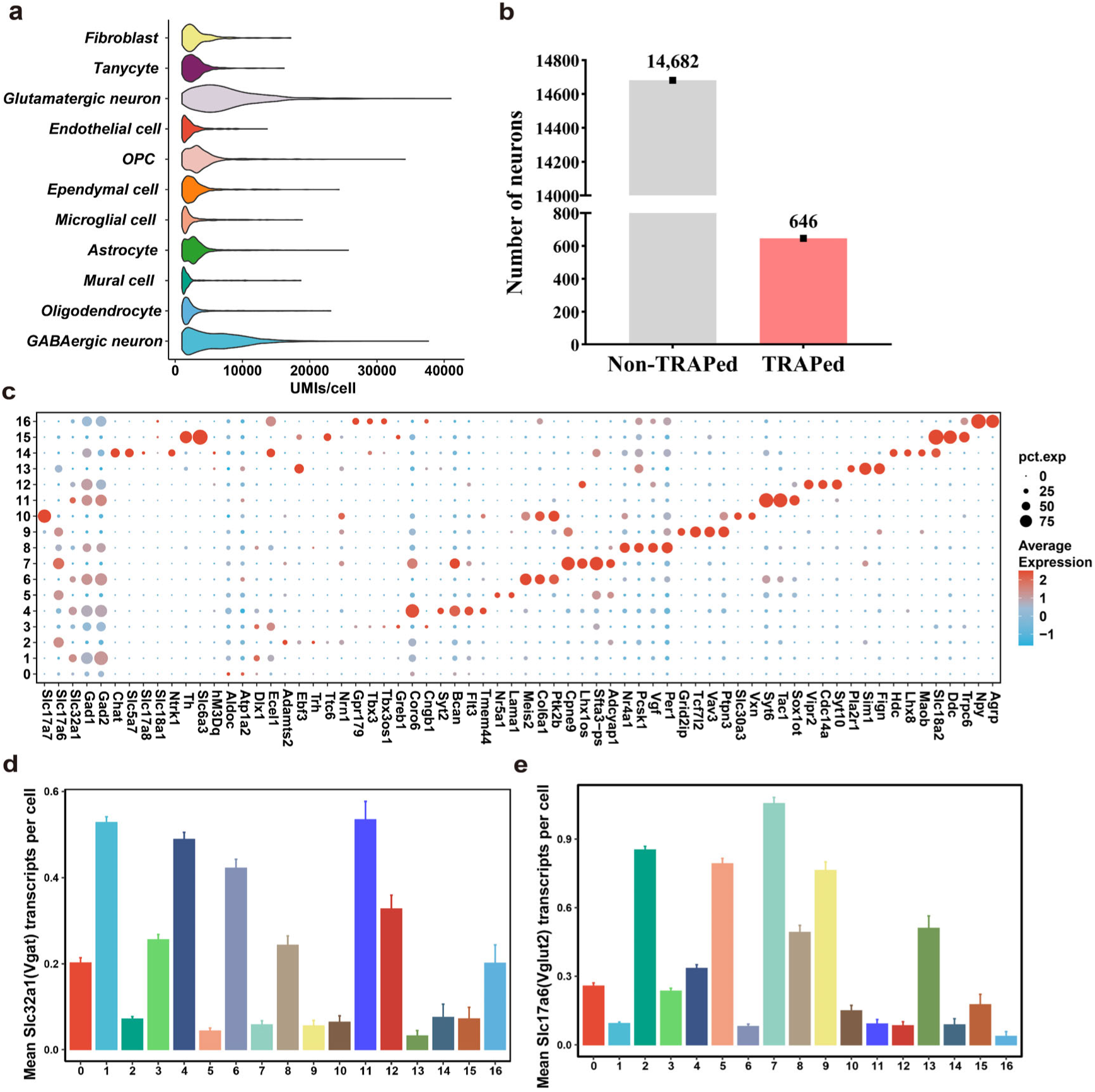
Cell classification and the gene expression characteristics in various clusters of neurons. **a,** Violin plot of the distribution of UMIs per cell for each main cell class (fibroblasts, n=190 cells, tancytes, n=488 cells, Glutamatergic neurons, n=1439 cells, endothelial cells, n=251 cells, OPCs, n=687 cells, ependymal cells, n=720 cells, microglia cells, n=670 cells, astrocytes, n=3,421 cells, mural cells, n=250 cells, oligodendrocytes, n=4,082 cells, and GABAergic neurons, n=13,889 cells). **b,** Quantification of the number of TRAPed cells. **c,** Dot Plot displays the expression of annotated markers within each cell cluster. Color intensity reflects the mean expression level, while the dot size corresponds to the percentage of cells expressing the gene in that cluster. **d, e,** Mean transcripts for Vgat **(d)** and Vglut2 **(e)** across various clusters of neurons.

**Fig. S4.**
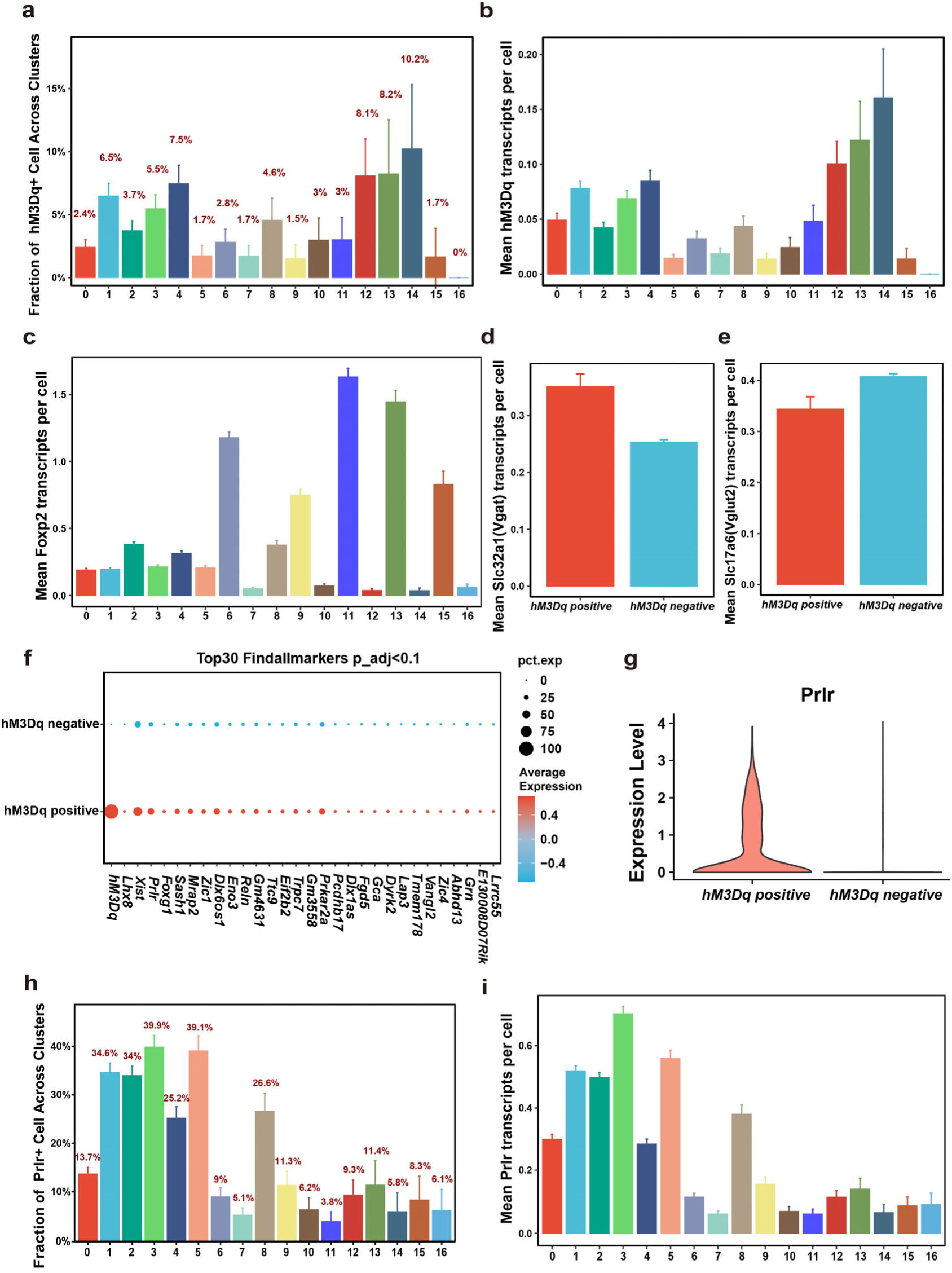
The expression of maker genes across neuronal clusters. **a, b,** Fraction of *hM3Dq*^+^ cells **(a)** and mean *hM3Dq* transcripts **(b)** across various clusters of neurons. **c,** Mean transcripts for *Foxp2* across various clusters of neurons. **d, e,** Mean transcripts for Vgat **(d)** and Vglut2 **(e)** across *hM3Dq*^+^ and *hM3Dq*^-^ neurons. **f,** Top30 markers between *hM3Dq^+^*and *hM3Dq^-^* groups. **g,** Expression level of *Prlr* between *hM3Dq*^+^ and *hM3Dq*^-^ groups. **h, i,** Fraction of *Prlr*^+^ cells **(h)** and mean *Prlr* transcripts **(i)** across *hM3Dq*^+^ and *hM3Dq*^-^ neurons.

**Fig. S5.**
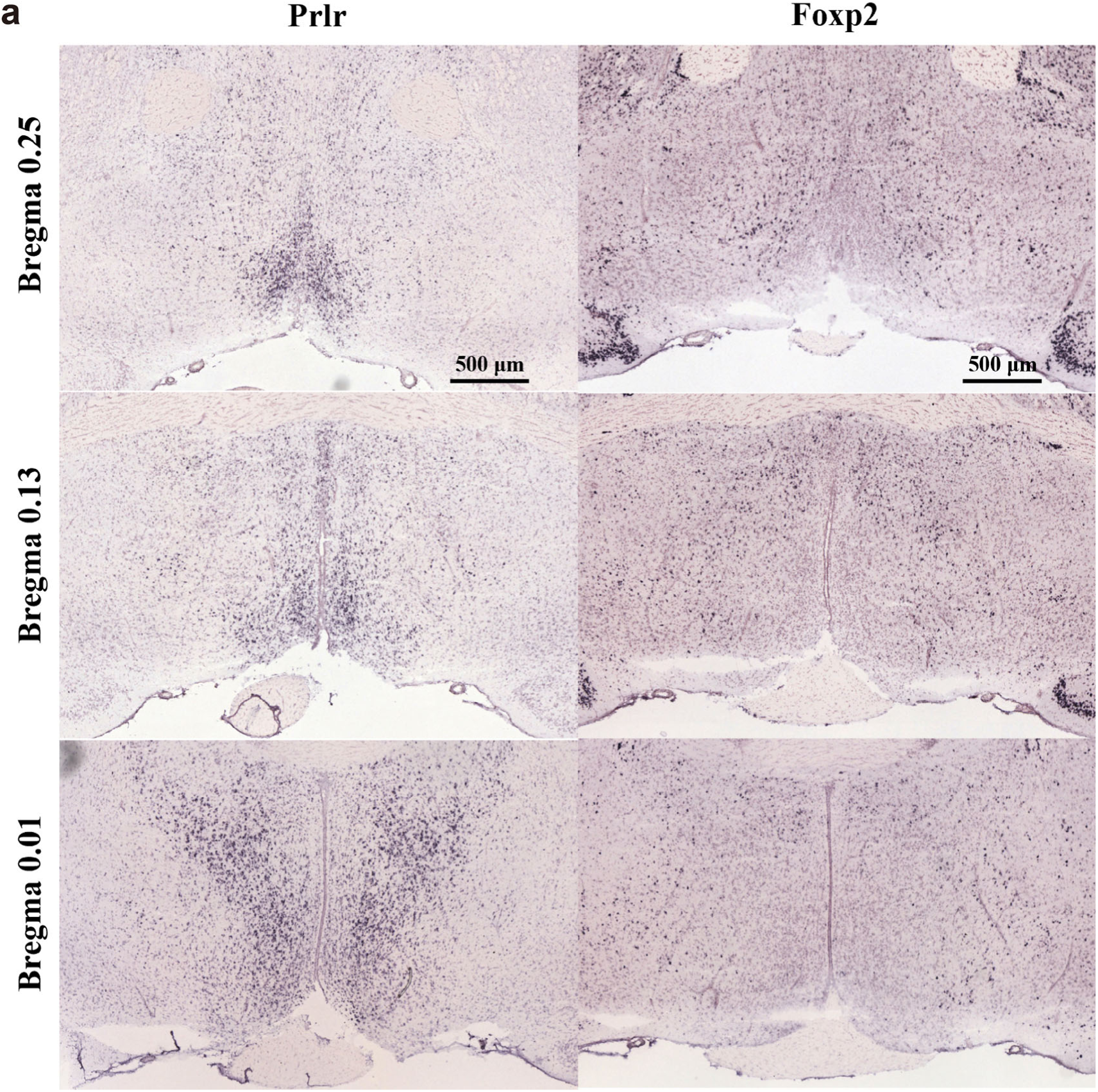
In situ hybridization results of gene expression in the Allen brain atlas for *Prlr* and *Foxp2*. Expression of *Prlr* (left) and *Foxp2* (right) in adult mouse brain. Allen Mouse Brain Atlas, *Prlr*, mouse.brain-map.org/gene/show/18879. *Foxp2*, mouse.brain-map.org/experiment/show/72079884.

**Fig. S6.**
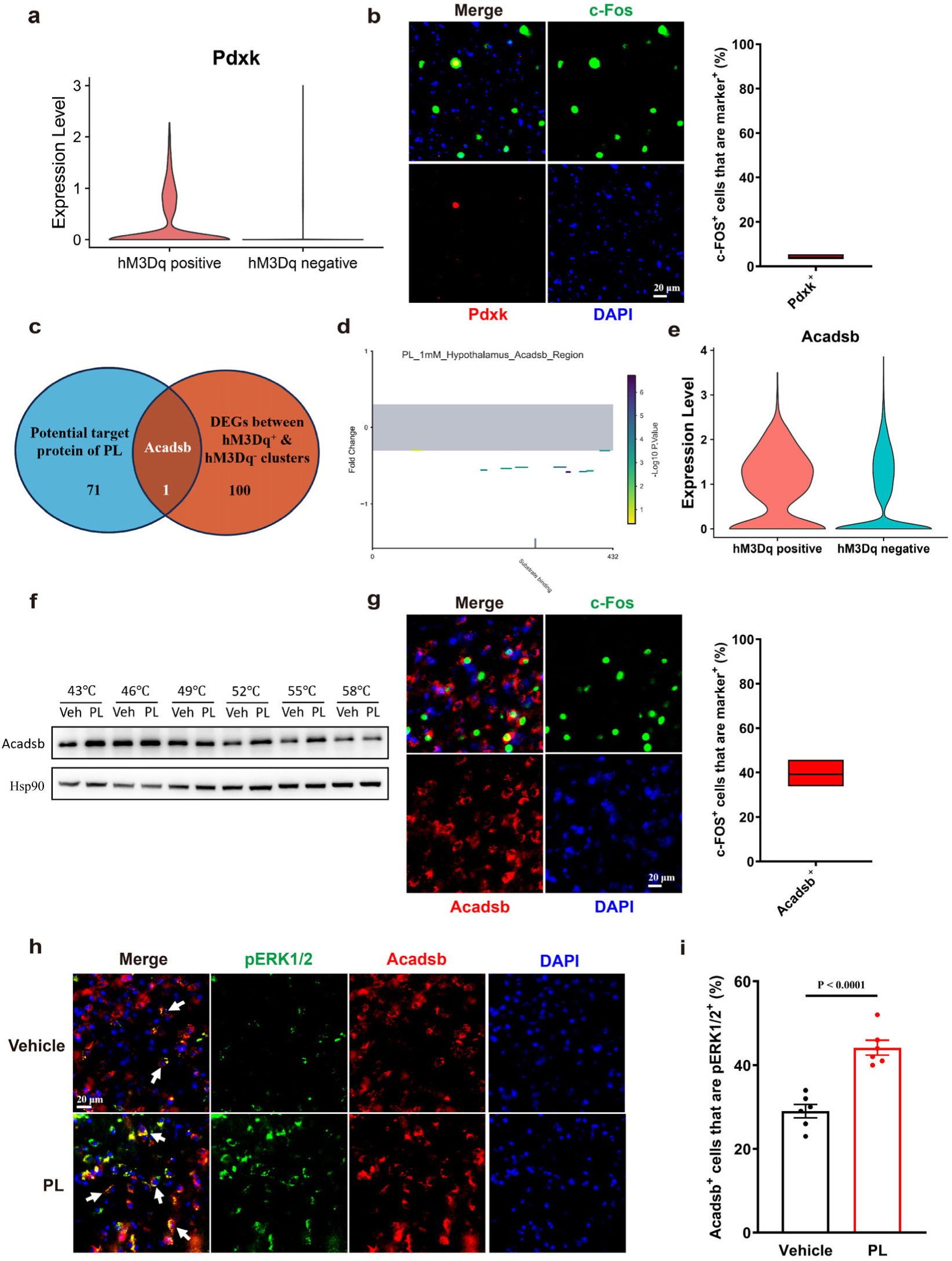
PL targets Acadsb to activate the ERK-FOS pathway. **a,** Expression level of *Pdxk* in hM3Dq positive cluster (*hM3Dq*^+^) and hM3Dq negative cluster (*hM3Dq*^-^). **b,** Pdxk and c-Fos expression in MAP in C57BL/6 mice intraperitoneal injection of P57 after 2 h with quantification (n=6 mice). **c,** Venn diagram shows the intersection of 26 potential targets of PL with the differentially expressed genes (DEGs) between *hM3Dq*^+^ cluster and *hM3Dq*^-^ cluster in the snRNA-seq data, with only one being Acadsb. **d,** PeptideMap of the changed peptide segments of Acadsb in mouse hypothalamic tissue after incubation with PL. **e,** Expression level of *Acadsb* in *hM3Dq*^+^ cluster and *hM3Dq*^-^ cluster. **f,** Protein thermal stability changes of Acadsb after PL (1 mM) treatment in HEK293T cells. **g,** Acadsb and c-Fos expression in MAP in C57BL/6 mice intraperitoneal injection of P57 after 2 h, with quantification (n=6 mice). **h, i,** Acadsb and pERK1/2 (T202/Y204) expression in MAP in C57BL/6 mice intraperitoneal injection of PL or vehicle 30 min later, white arrow indicates Acadsb^+^ and pERK1/2^+^ cells **(h)**, with quantification **(i)** (n=6 mice). Student’s t test used was two-sided. Data are presented as mean values ± s.e.m.

**Fig. S7.**
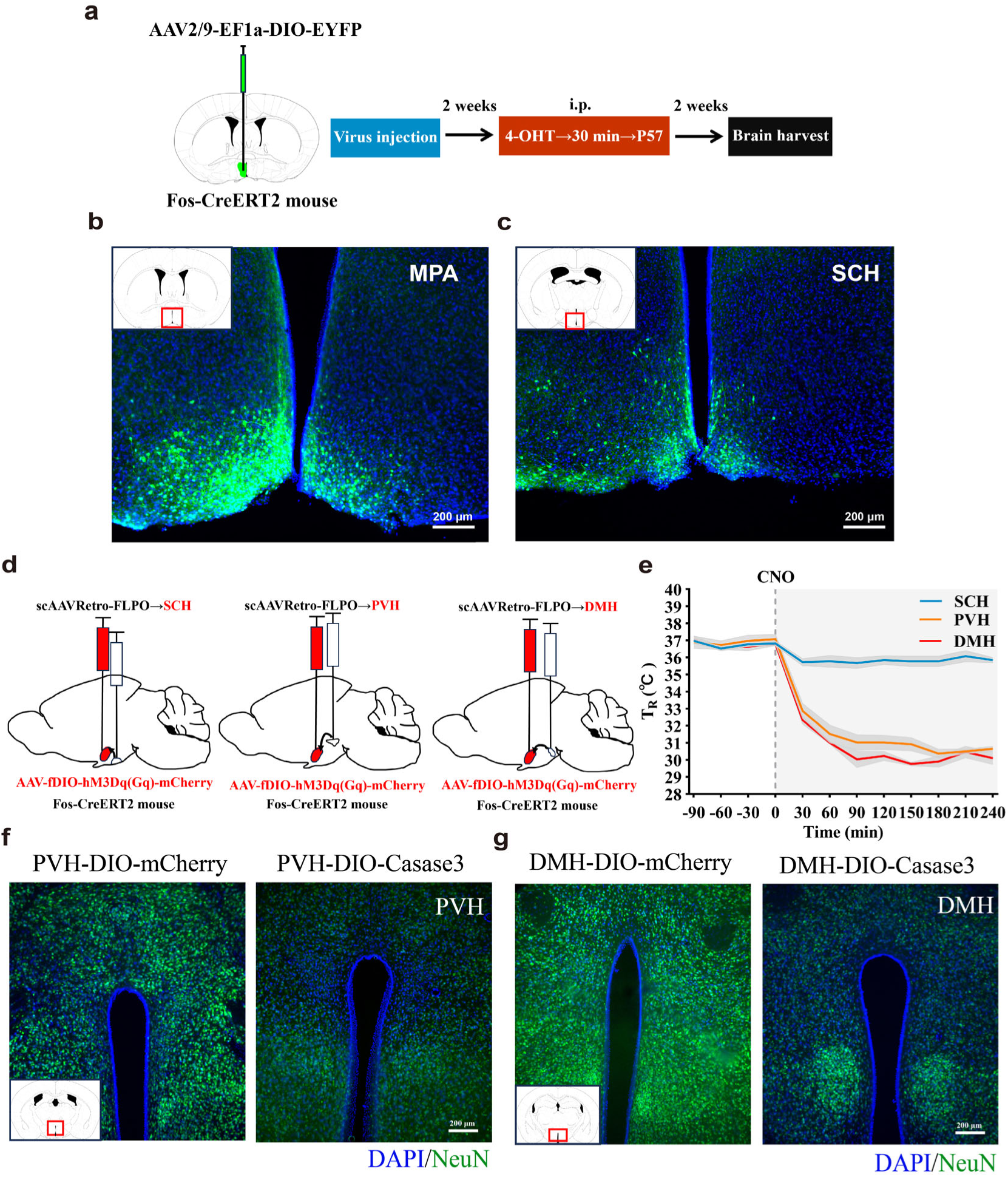
MPA-PVH circuit mediates P57-induced hypothermia. **a,** Schematic shows the process of using anterograde tracer viruses to discover downstream projections of neurons activated by P57 in MPA in *Fos-CreERT2* mice. **b,** Expression of the AAV2/9-EF1a-DIO-EYFP in MPA in *Fos-CreERT2* mice (n=5 mice). **c,** Representative images of projections to SCH. **d,** Schematic shows chemogenetic circuit manipulation of three downstream brain regions of the MPA by retrovirus in combination with the FLPO-fDIO system. **e,** T_R_ of *Fos-CreERT2* mice in **(d)** before and after intraperitoneal injection of CNO (n=4 mice in SCH group, 4 mice in PVH group and 3 mice in DMH group). The dashed line indicates the onset of CNO administration, shading indicates error bars. **f, g,** Representative images of neuronal death in PVH **(f)** and DMH **(g)**.

**Fig. S8.**
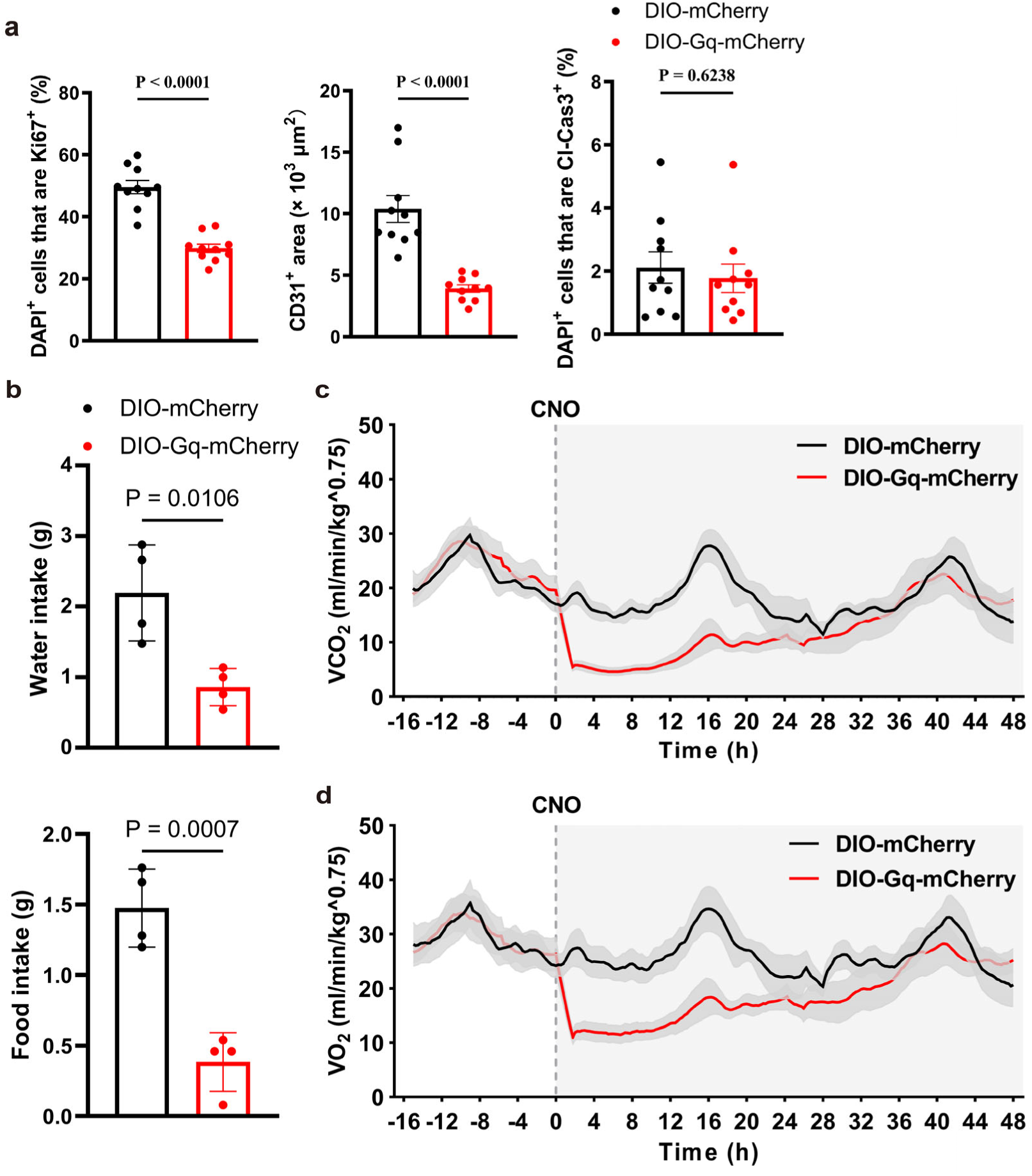
Long-term low temperature reduces energy metabolism. **a,** Quantification of immunofluorescence staining of Ki67, CD31 and Cl-Cas3 in tumor tissue (n=10 random fields per group). **b,** Water intake (top) and food intake (bottom) of *Foxp2-Cre* mice bearing tumor (n=4 mice in each group) in metabolic chambers over 24 hours after CNO administration. **c, d,** Volume of Carbon dioxide (VCO_2_) **(c)** and volume of oxygen (VO_2_) **(d)** of *Foxp2-Cre* mice bearing tumor from day 7 to day 9 (n=4 mice in each group). Student’s t test used was two-sided. Data are presented as mean values ± s.e.m.

**Fig. S9.**
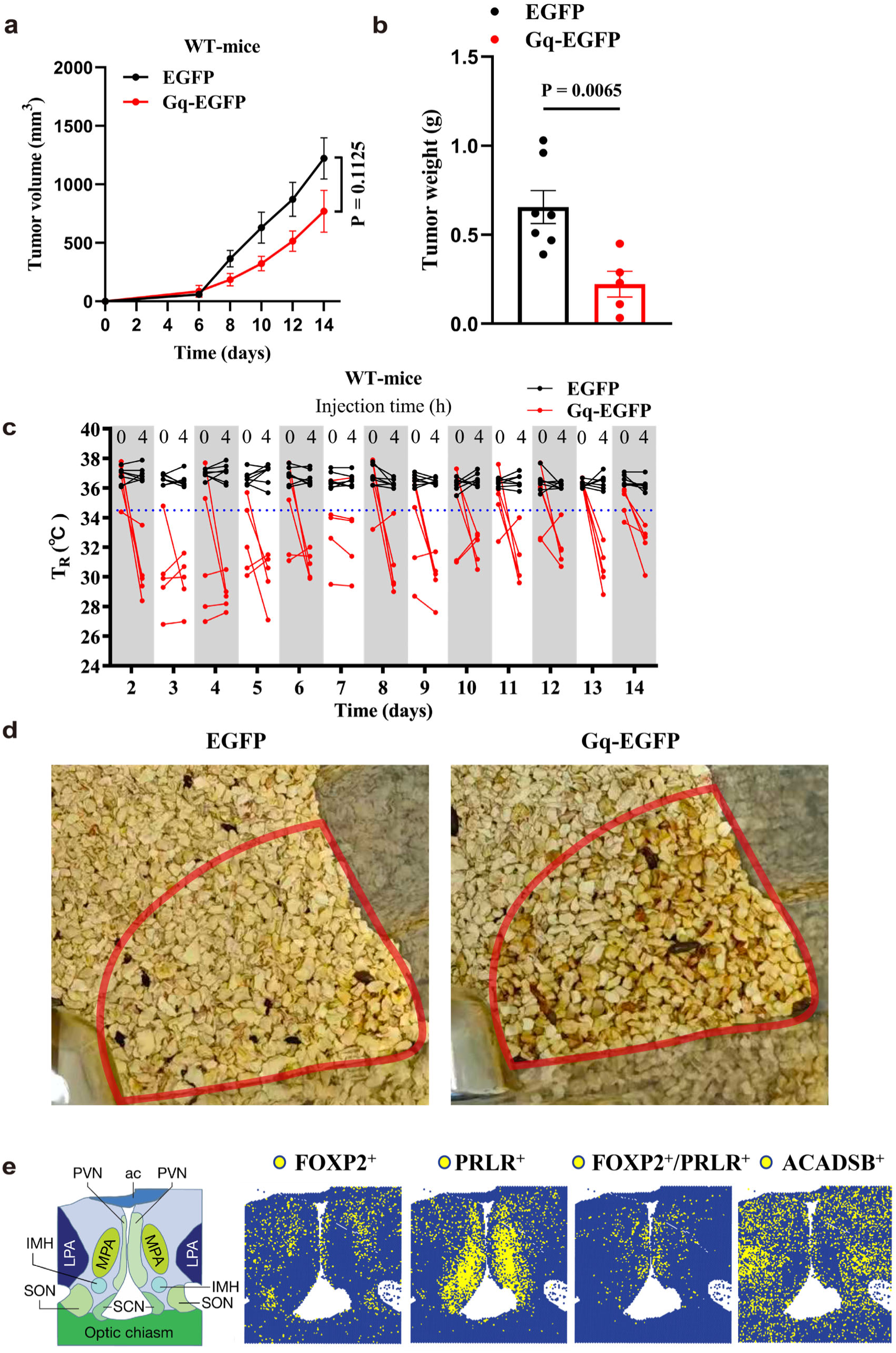
Neuronal control of therapeutic hypothermia inhibits tumor growth. **a, b,** Tumor volume **(a)** and tumor weight in day 14 **(b)** of C57BL/6 mice (WT mice) that received virus injection (n=5 mice in Gq-EGFP group and 7 mice in EGFP group). **c,** T_R_ of mice in **(a)** during experiment period. The administration and T_R_ measurement of mice are described in detail in the methods section. **d,** A representative image showing that the mice in the Gq-EGFP group had an increased urine output, causing the corn cob bedding to become wet. **e.** Expression analysis of *FOXP2, PRLR, FOXP2/PRLR and ACADSB* in human hypothalamic spatial transcriptome data. The reference atlas figure showed on the left^33^. Student’s t test used was two-sided. Data are presented as mean values ± s.e.m.

